# DR-SIP: Protocols for Higher Order Structure Modeling with Distance Restraints- and Cyclic Symmetry-Imposed Packing

**DOI:** 10.1101/500397

**Authors:** Justin Chan, Jinhao Zou, Chi-Hong Chang Chien, Rong-Long Pan, Lee-Wei Yang

## Abstract

**Motivation:** Quaternary structure determination for proteins is difficult especially for transmembrane proteins. Even if the monomeric constituents of complexes have been experimentally resolved, computational prediction of quaternary structures is a challenging task particularly for higher order complexes. It is essential to have a reliable computational protocol to predict quaternary structures of both transmembrane and soluble proteins leveraging experimentally determined distance restraints and/or cyclic symmetry (C*_n_* symmetry) found in most homo-oligomeric transmembrane proteins.

**Results:** We survey 115 X-ray crystallographically solved structures of homo-oligomeric transmembrane proteins (HoTPs) to discover that 90% of them are C*_n_* symmetric. Given the prevalence of C*_n_* symmetric HoTPs and the benefits of incorporating geometry restraints in aiding quaternary structure determination, we introduce two new filters, the distance-restraints (DR) filter and the Symmetry-Imposed Packing (SIP) filter which takes advantage of the statistically derived tilt angle cutoff and the C_n_ symmetry of HoTPs without prior knowledge of the number (“*n*”) of monomers. Using only the geometrical filter, SIP, near-native poses of the 115 HoTPs can be correctly identified in the top-5 for 52% of all cases, or 49% among the HoTPs having an *n* >2 (~60% of the dataset), while ZDOCK alone returns 41% and 24%, respectively. Applying only SIP to three HoTPs with distance restraints, the near-native poses for two HoTPs are ranked 1^st^ and the other 7^th^ among 54,000 possible decoys. With both filters, the two remain 1^st^ while the other improved to 2^nd^. While a soluble system with distance restraints is recovered at the 1^st^-ranked pose by applying only DR.

**Availability and Implementation:** https://github.com/capslockwizard/drsip

**Supplementary information:** Supplementary methods and results are available.

## INTRODUCTION

The quaternary structure of proteins provides atomistic details which can be used to study the mechanisms underlying the function of these proteins. A large subset of these protein complexes are transmembrane proteins which constitute 20-30% of the proteome (Fagerberg *et al*., 2010) and are the second most common drug targets (Rask-Andersen *et al*., 2014), after enzymes. A survey on RCSB PDB (Rose *et al*., 2017) shows that about 66% of α-helical transmembrane proteins are homo-oligomeric transmembrane proteins (HoTPs), while 92% of HoTPs are cyclic (Cn) symmetric (Fig. 1).

**Fig. 1.**
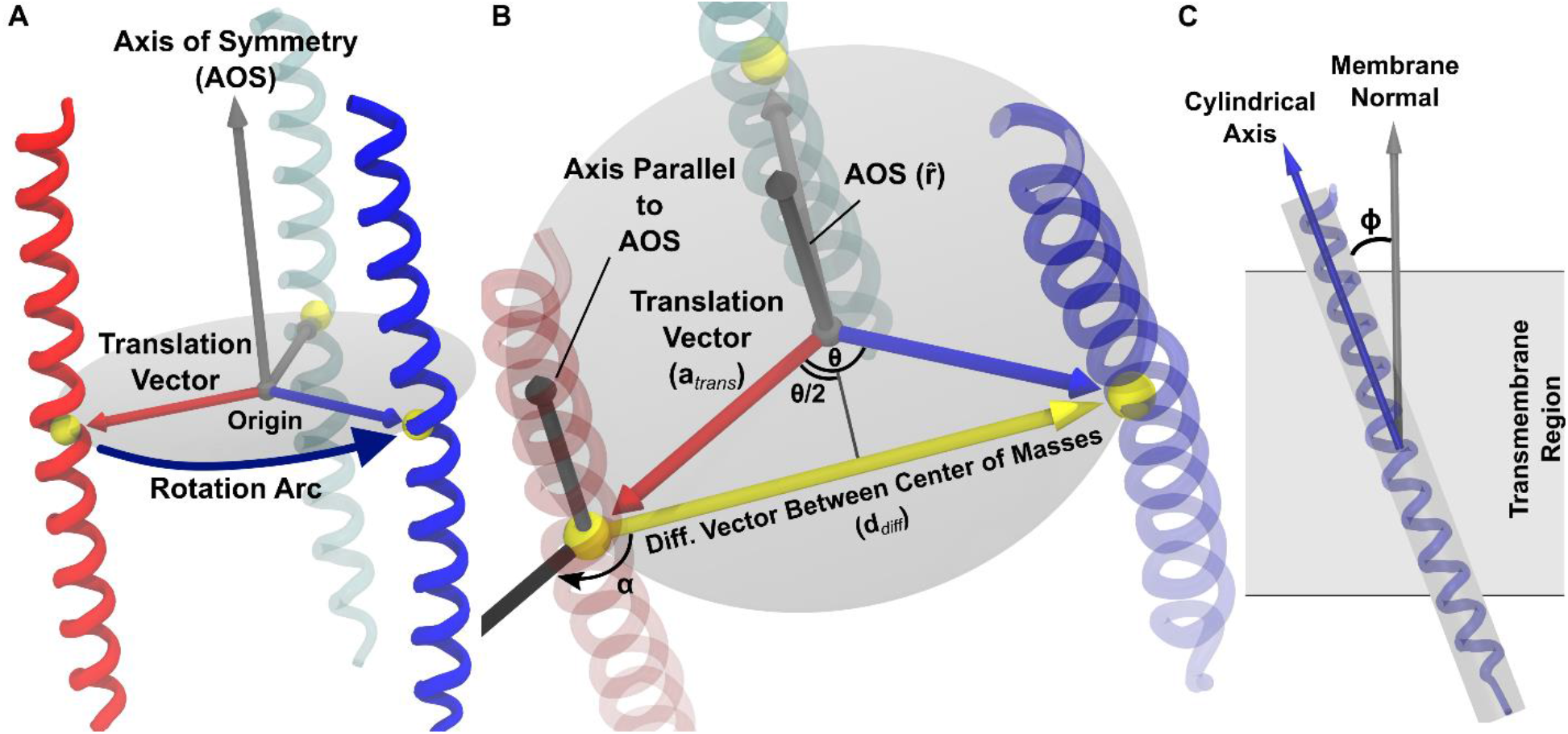
C_n_ symmetry and tilt angle of HoTPs. (**A**) Three helices with C3 symmetry about the axis of rotation/symmetry (AOS). (**B**) The translation vector (**a***_trans_*) can be expressed as a function of angle of rotation (θ) and difference vector between the centers of mass of the two monomers (**d***_diff_*) such that **a***_trans_* is obtained by rotating **d***_diff_* by α = 90° + θ/2 degree about the AOS (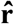) and then rescaled to ||**d***_diff_*||(2sin(θ/2)) for its magnitude. (**C**) Tilt angle (ɸ) is the acute angle between the cylindrical axis and membrane normal (assumed to be parallel to AOS).

Experimentally solving the ternary and quaternary structures of transmembrane proteins is challenging. The general difficulty is in purifying proteins in sufficient quantity and purity. For X-ray crystallography, a major challenge after obtaining proteins is in screening for the optimal conditions to reconstitute and crystalize the proteins (Moraes *et al*., 2014). On the other hand, solution NMR is limited to protein complexes with a high tumbling rate (Liang and Tamm, 2016) which limits the size of the complex while solid-state NMR is limited by spectral crowding, which is further aggravated by large complexes (Liang and Tamm, 2016).

Computational methods such as homology docking has been applied to predict quaternary structures which are evolutionarily conserved (Levy *et al*., 2008). These methods (Szilagyi and Zhang, 2014; Bertoni *et al*., 2017) identify structurally solved homologous protein complexes, not just constituent proteins, as the templates for the homology modelling. However, to predict higher order structures for proteins that lack any structural template of homologs, physiochemically driven molecular docking could be the only option to go about.

Molecular docking is a method that usually generates tens of thousands of orientations (poses) of two proteins bound to one another which are then rank-ordered by a scoring function that evaluates the fitness (or “energy”) of these poses (Huang, 2014). Identifying near-native poses among the >10,000 generated poses is difficult because of the efficient sampling in molecular docking comes at the expense of accurately evaluating the solvent and entropy’s energy (Brooijmans and Kuntz, 2003). Common docking packages include ZDOCK (Pierce *et al*., 2011), ClusPro (Kozakov *et al*., 2017), GRAMM-X (Tovchigrechko and Vakser, 2006), RosettaDock (Chaudhury and Gray, 2008) and HADDOCK (van Zundert *et al*., 2016), where the latter two allow for experimentally determined restraints or prior knowledge of pairwise distances to constraint and filter the docking results.

Predicting interactions between membrane proteins, DOCK/PIERR (Viswanath *et al*., 2015), ROSETTA:MPdock (Alford *et al*., 2015) and Memdock (Hurwitz *et al*., 2016) generate poses whose transmembrane helices in the constituent monomers are embedded in the membrane bilayer, which enriches the near-native poses among the decoys. There are also docking packages that specifically perform C_n_ symmetric docking such as SymmDock (Schneidman-Duhovny *et al*., 2005), ClusPro (Kozakov *et al*., 2017), M-ZDOCK (Pierce *et al*., 2005), ROSETTA:MPsymdock (Alford *et al*., 2015) and HADDOCK (van Zundert *et al*., 2016). Although these methods are successful at predicting the native complexes, they require prior knowledge of the number of constituent monomers in the complex, or the “order of symmetry”.

Integrative approaches, combining multiple sources of experimental data with computational methods, have been used to confine the search space during generation of poses or to filter out the poses that are incompatible with the data thereby enriching the near-native ones among the docking decoys (Alber *et al*., 2008). Biochemically and biophysically determined distance restraints include but are not limited to the data obtained from mutagenesis, cross-linking, hydrogen/deuterium exchange, electron microscopy (Mitra and Frank, 2006), small-angle X-ray scattering (Yamagata and Tainer, 2007), NMR chemical shift perturbations (Gupta *et al*., 2013; Chang *et al*., 2016; Khan *et al*., 2018) and single molecule resonance energy transfer (smFRET) (Dimura *et al*., 2016; Choi *et al*., 2010; Muschielok *et al*., 2008), which have been used to predict quaternary structures (Alber *et al*., 2008).

Among these biophysical methods, smFRET measures the distance between two fluorophore dyes acting as labels attached to residues in proteins or nucleotides to study protein folding, protein-protein interaction and protein-DNA/RNA interaction (Sasmal *et al*., 2016). However, these distances suffer from uncertainties due to the effect of the local environment on the dye pairs (Dimura *et al*., 2016; Kalinin *et al*., 2012; Muschielok *et al*., 2008).

In this study, we introduce two new docking protocols including the two new filters, the Distance Restraints (DR) filter and Symmetry-Imposed Packing (SIP) filter, to facilitate quaternary structure determination for soluble and transmembrane proteins. Here the “DR” filter requires experimentally measured distance restraints but the geometric filter, SIP, requires no experimental input. The SIP filter removes the docking poses (homo-dimer) that deviate significantly from its ideal C_n_ symmetry without prior knowledge of the order of symmetry (the *n* in C_n_) and those having large tilt angles. Given the uncertainties of the absolute distances obtained from smFRET, the DR filter uses the agreement in the relative ranking of these distances by correlating the smFRET-measured intramolecular distances and their counterparts in the docking poses. The cutoffs for these filters are derived from the statistical analyses of a dataset containing the X-ray structures of 118 α-helical HoTPs (see **Results**). Evaluating only the SIP filter without distance restraints on all 115 C_n_ symmetric HoTPs from the dataset shows the near-native poses in 64% of HoTPs can be recovered within the top-10 results starting from the 54,000 poses generated by ZDOCK, as compared to the 45% recovered by ZDOCK alone. For the 59% of HoTPs that are larger than dimers (*n*>2), SIP recovers 57% of the native poses in the top-10 while ZDOCK alone merely recovers 28%. Furthermore, for four systems (soluble: Syt1-SNARE complex, three HoTPs: MscL, *Vr*H+-PPase and *Ct*H+-PPase) with experimentally determined distance restraints, the DR-SIP filters return a near-native pose in the top rank for three systems and 2^nd^-rank for MscL. Without the DR filter, a near-native pose for MscL is 7^th^-ranked while the other two HoTPs are still top-ranked. Other than ZDOCK, DR-SIP is applied to GRAMM-X’s results and the near-native poses are enriched up to ~167-fold such that they make up one out of every three remaining poses.

## METHODS

### Docking Protocols

DR-SIP contains two docking protocols (Fig. 2). The first protocol is for predicting the quaternary structures of HoTPs (dashed line in Fig. 2) and the other for soluble proteins (dotted line in Fig. 2). The docking protocols make use of the DR and SIP filters. The SIP filter (see below and **Supplementary Methods**) consists of the C_n_ symmetry root-mean squared deviation (RMSD) and the tilt angle criteria.

**Fig. 2.**
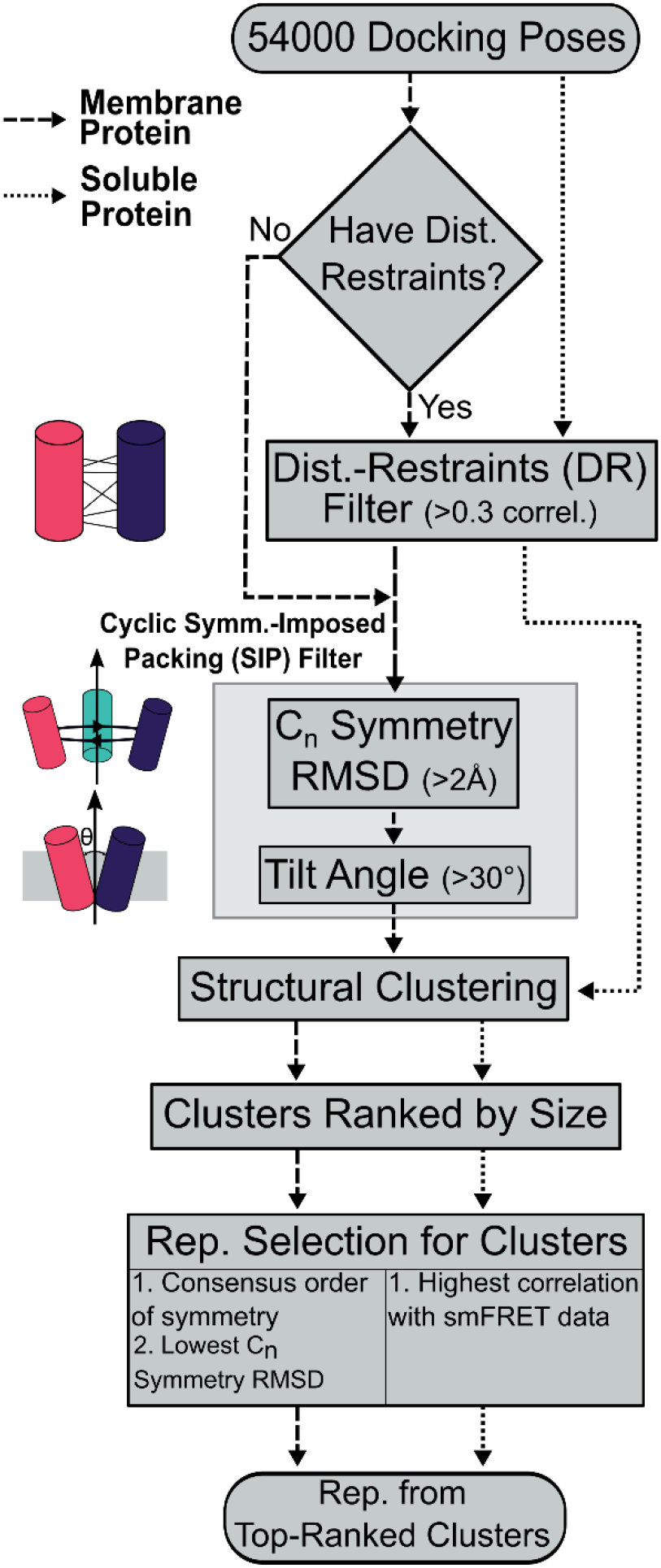
Flowchart of DR-SIP’s membrane and soluble protein docking protocols. Each cluster is represented by one member of the cluster. The final results are the representatives ranked based on the size of the clusters.

For every system, 54,000 docking poses are generated using ZDOCK 3.0.2 (Pierce *et al*., 2011) with 6° rotational sampling (dense sampling).

C_n_ symmetry root-mean-squared deviation (C_n_ RMSD) measures the current pose’s deviation from its closest ideal C_n_ symmetric pose (see Methods). The axis and order of symmetry (*n*) are estimated from each docking pose containing two monomers and used to generate the ideal C_n_ symmetric pose. Based on our statistical analysis (see Figs. 3C and 4), non-Cn symmetric poses with >2Å C_n_ RMSD are removed.

**Fig. 3.**
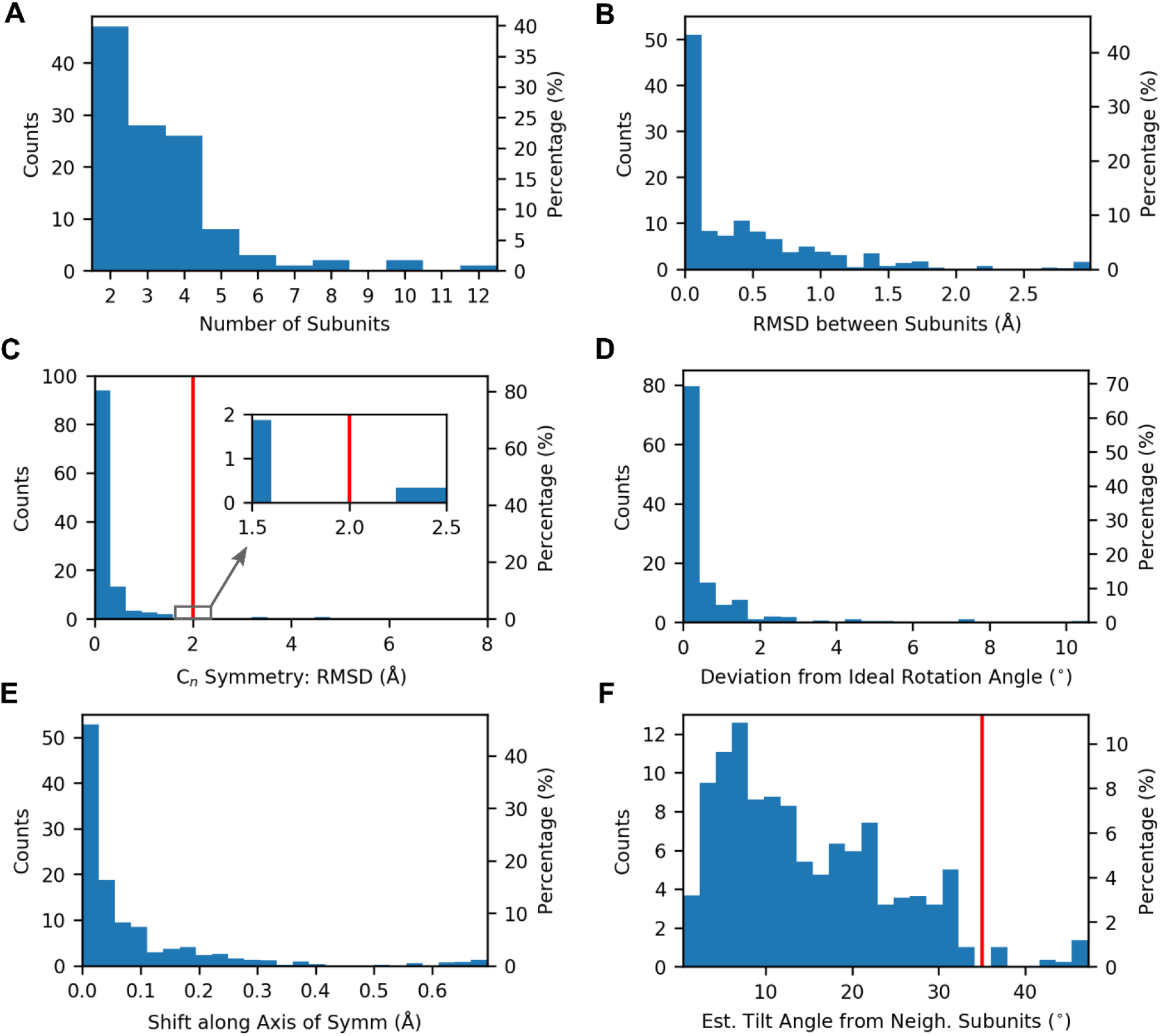
Statistical distributions of the 118 HoTPs with parallel orientation. The left axis shows the counts of each bin while their percentages (out of 115 or 118 structures) are shown on the right axis. (**A**) Distribution of oligomeric states of the HoTPs reveals 85.6% (101/118) of the structures are dimers, trimers and tetramers. (**B**) Conformational differences between neighboring monomers measured by root-mean squared deviation (RMSD) of Cα atoms, shows that the conformational differences are small, all are <3Å while 87.0% <1Å. (**C**) The C_n_ symmetry RMSD measures the translational and rotational differences for a pair of neighboring monomers caused by deviation from ideal C_n_ symmetry (see Methods). The distribution includes the HoTPs within 8Å (except for PDB ID: 5HK1). The cutoff of <2Å (see red line and inset) is used to identify C_n_ symmetric HoTPs. 115 (97.5%) out of the 118 structures are within this cutoff. The remaining three structures (**Supplementary Results**), PDB IDs: 2GIF, 2QTS and 5HK1, with the lowest RMSD between neighboring monomers of 2.5Å, 4.6Å, 21.1Å, respectively, were classified as non-Cn symmetric. (D and E) The deviation from ideal C_n_ symmetry for the remaining 115 structures can be decomposed into *rotational differences* and *translational differences* which are measured by the (**D**) deviation from ideal rotation angle and (**E**) shift along (parallel to) the AOS, respectively. The shift here refers to the amount of the translational difference between the two monomer’s COM (**d**_*diff*_, Fig. 1B) that are parallel to the AOS. (**F**) The tilt angle distribution shows the acute angle between the direction that the monomers span the membrane and the membrane normal (AOS). 97.4% of the 115 C_n_ symmetric structures have tilt angles <35° (red line) which is the cutoff employed in the SIP filter. (**B-F**) are calculated using all neighboring pairs of monomers in each HoTP. Each pair is weighted to contribute 1/*n* to the counts except for homo-dimers where the weight is one.

**Fig. 4.**
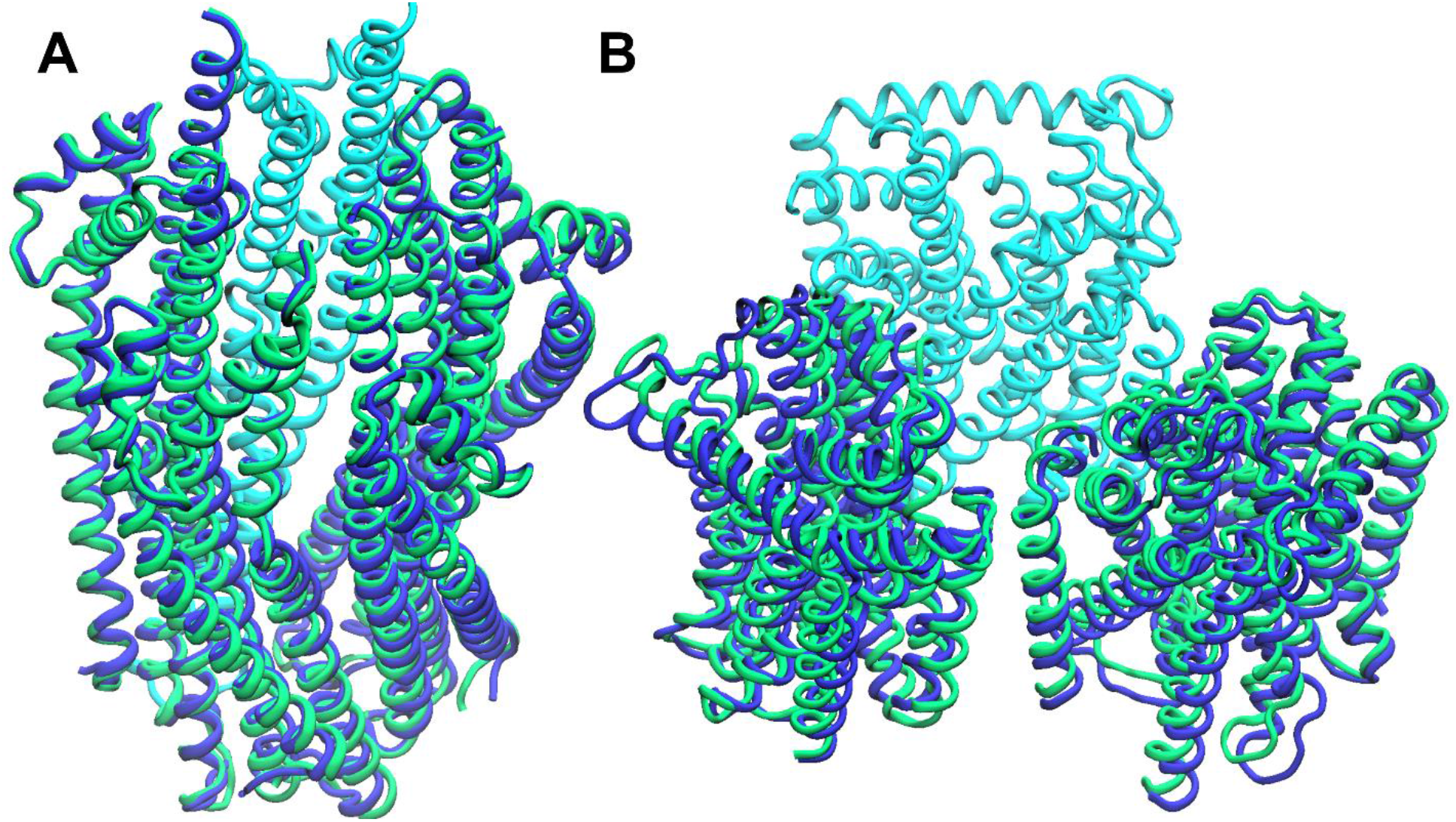
Comparison of the X-ray structure (blue) and the ideal C_n_ symmetric complex (green) for the two HoTPs closest to the C_n_ symmetry RMSD cutoff of 2Å. The complexes are superimposed at chain A (cyan). (**A**) The protein that is closest to and below the cutoff is the ExbB/ExbD complex (PDB ID: 5SV0, C_n_ symmetry RMSD: 1.6 Å). (**B**) The protein that is closest to, but above the cutoff is the multidrug efflux pump subunit AcrB (PDB ID: 2GIF, RMSD: 2.5Å). There are more visible differences when the RMSD is above the chosen cutoff of 2Å.

While, tilt angle is the acute angle between the membrane’s normal and the direction the monomer spans the membrane (Fig. 1C). Assuming the membrane normal is the axis of symmetry (AOS) of HoTPs, poses having tilt angles >35° are removed in accordance with the observed HoTP tilt angle distribution (see Fig. 3F).

The distance-restraints (DR) filter removes non-native poses that are incompatible with the experimentally measured constraints (herein smFRET-characterized distances) where relative distances rather than absolute distances are used. This is measured by the Pearson and Spearman correlations between the smFRET-measured distances and the distances of the corresponding labeled residues (between Cα atoms) in the docking pose. Poses are removed if they have ≤0.3 correlation.

After applying the filters, the remaining poses are clustered using agglomerative clustering with the average linkage criteria (Manning *et al*., 2008), see **Supplementary Methods** for details. RMSD is used to measure the difference/distance between all pairs of poses. A cutoff of 12Å is employed and clusters with fewer than three poses are removed. The clusters for proteins are ranked based on their sizes in descending order.

For HoTPs, each cluster is assigned a consensus order of symmetry (*n*), such that most members in the cluster have this ‘*n*’. The representative pose of each cluster must have the consensus order of symmetry and is the closest to the ideal C_n_ symmetry as measured by the smallest C_n_ symmetry RMSD.

For soluble proteins, the representative pose of each cluster has the highest Spearman correlation with the distance restraints data (smFRET data). If there are ≥2 poses with the same Spearman correlation, the one with the higher Pearson correlation is chosen as the representative.

The final results returned to the user are the representative poses from the top clusters. Details of the filters and clustering method are in the **Supplementary Methods**.

### Extracting C_n_ Symmetry Parameters from an Ideal C_n_ Symmetric Pose

As explained before, also true for this and the next sections, each ideal C_n_ symmetric HoTP pose contains two identical and neighboring monomers (the blue and red monomers of the C3 complex in Fig. 1A) that is a part of (C_n>2_) or is the full (C_2_) protein complex. The number of monomers in a complex can be determined by dividing 2π with the angle of rotation between these two monomers (see below).

The two monomers, A and A’, are related by

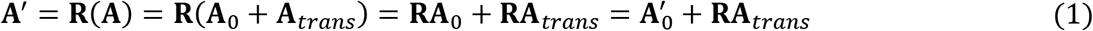

where **A** and **A’** are *3xM* coordinate matrices of *M* atoms for A and A’, respectively. R is a rotation matrix that rotates about the AOS by 2π/*n* rad (*n* is the order of symmetry). In Equation (1), the COM of the n-mer complex is assumed to be at the origin (e.g. the center of the trimer in Fig 1A). When the COM of A and A’ are placed at the origin, their coordinate matrices are **A**_0_ and **A’**_0_, respectively. Let **A***_trans_* = **a***_trans_* [1, 1,…1]_1×M_ where **a***_trans_* is the 3×1 vector pointing from the origin to the COM of A.

The docking pose provides **A** and **A’** which are used to compute the AOS (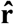), angle of rotation (*θ_r_*), and the translation vector (**a***_trans_*). These three parameters define the C_n_ symmetric relationship between **A** and **A’** which can be used to reconstruct the n-mer complex.

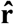 and *θ_r_* can be obtained from **R**. As for **R**, from Equation (1), **A’**_0_ = **RA**_0_, the Kabsch algorithm is used to find an **R** that minimizes the RMSD between **A’**_0_ and **RA**_0_ (Kabsch, 1976; Cock *et al*., 2009).

Given **R**, 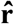 is the eigenvector associated with the unity eigenvalue such that 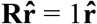, because the AOS is invariant when operated on by **R**.

As for *θ_r_*, it can be found by extracting cos *θ_r_* and sin *θ_r_* from **R** which can be formulated as (Jia, 2017):

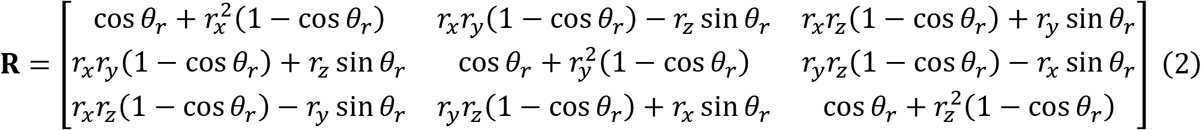

where *θ_r_* is the rotation angle and 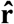 (*r_x_, r_y_, r_z_*)is the AOS (unit vector). The trace of **R** is 1 + 2cos *θ_r_*. Thus cos *θ_r_* can be obtained.

While, sin *θ_r_* is obtained as follows

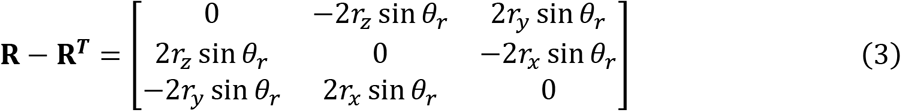

where 2*r_x_*sin *θ_r_* + 2*r_y_*sin *θ_r_* + 2*r_z_*sin *θ_r_* = 2sin *θ_r_* (*r_x_* + *r_y_* + *r_z_*). With 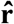, we solve for sin *θ_r_*.

Finally, *θ_r_* = sign(sin *θ_r_*) arccos *θ_r_*. Alternatively, *θ_r_* can be computed using the atan2 function which is implemented in most mathematical software libraries (van der Walt *et al*., 2011).

**a***_trans_* is obtained by rotating **d***_diff_* about 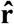 by α radians (Fig. 1B), which is then rescaled to ||**a***_trans_*|| based on its trigonometric relationship with **d***_diff_*

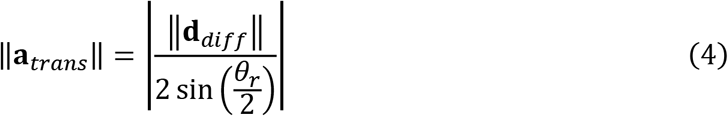

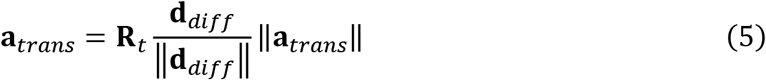

where **d***_diff_* is the vector from the COM of **A** to **A**’ and **R***_t_* is a rotation matrix, rotating α = [-sign(*θr*) (|*θ_r_*|/2 + π/2)] rad about 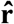 (Fig. 1B).

### Constructing the Ideal C_n_ Symmetric Pose that is Closest to a Non-Ideal Docking Pose or X-Ray Crystallographically Solved Structure

No poses obtained from docking or experimentally resolved structures are in perfect C_n_ symmetry (non-ideal). Our goal here is to obtain an ideal C_n_ symmetric pose that is closest to the non-ideal pose. The deviation of the non-ideal pose from the ideal pose is used to determine if the non-ideal pose is C_n_ symmetric (see C_n_ Symmetry RMSD in Supplementary Methods).

To construct the ideal C_n_ symmetric pose (A’*) from a non-ideal pose, the angle of rotation and translation vector are computed as follows

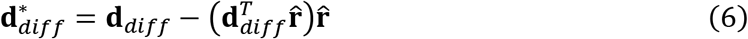

where *n* = [|2π/*θ_r_*| rounded to the nearest integer] (such that *θ_n_* = 2π / *n*) and 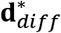 is the vector component of **d***_diff_* that is orthogonal to 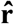. Note that, 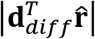 is the shift along the AOS (see **Supplementary Methods**).

Then, **a***_trans_* is computed with 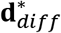 and θ*_n_* (see Equation (4) and (5)). To obtain the A’* pose, **A**_0_, **a***_trans_* and **R**\* are substituted into Equation (1), where the rotation matrix **R**\* rotates about 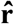 by *θ_n_*.

### Data Sets

#### Non-redundant X-ray structures of α-helical HoTPs

The PDB IDs of α-helical transmembrane proteins were taken from the mpstruct database (http://blanco.biomol.uci.edu/mpstruc/) and filtered with RCSB PDB's advanced filter (Rose *et al*., 2017) to remove redundant proteins and select for high-resolution X-ray structures (see Supplementary Methods). The structures were manually verified to be transmembrane and homo-oligomeric proteins resulting in the final data set containing 129 PDB IDs.

#### smFRET and structural data used for validation

The Syt1-SNARE smFRET data are obtained from Choi *et al*. (Choi *et al*., 2010) and filtered such that each residue pair’s transfer efficiency histogram has a single peak (Supplementary Fig. S1, Table S1 and S2). Syt1-SNARE’s reference structure is PDB ID: 5CCG (Zhou *et al*., 2015) but the C2A and C2B domains of Syt1 are from PDB ID: 2R83 (Huang *et al*., 2013) which were used to fix the missing residue GLU130 (see Supplementary Methods). For MscL, the smFRET data (Supplementary Fig. S2 and Table S3) for the closed state *E. coli* MscL are from Wang *et al*. (Wang *et al*., 2014) while the reference structure is PDB ID: 2OAR (Steinbacher *et al*., 2007). The smFRET data (Supplementary Table S4) for the *Ct*H^+^-PPase system are from Huang *et al*. (Huang *et al*., 2013) and the reference structure is predicted by homology modeling using SwissModel (Biasini *et al*., 2014) with the template structure from the *Vr*H^+^-PPase system sharing a sequence identity of 45.1% (PDB ID: 4A01 (Lin *et al*., 2012); the smFRET data for *Vr*H^+^-PPase are mapped from *Ct*H^+^-PPase to *Vr*H^+^-PPase based on the pairwise sequence alignment (**Supplementary Fig. S3**). Further details can be found in the Supplementary Methods.

### Implementation of DR-SIP

The DR-SIP package is implemented in Python 2.7 (McKinney, 2010; Gowers *et al*., 2016; Lam *et al*., 2015; Cock *et al*., 2009; Millman and Aivazis, 2011) and contains four modules: drsip, drsip-common, zdock-parser and docking-eval. The drsip module implements the docking protocols, filters and a command-line-interface (CLI) for users to run the DR-SIP docking protocols. When the docking results are derived from ZDOCK, the zdock-parser module is available to parse the ZDOCK output file, generate the docking poses and write the coordinates out to PDB files. On the other hand, the docking-eval module implements the CAPRI criteria used to evaluate the quality of docking poses. Lastly, drsip-common contains functions that are commonly used by the other modules.

Regular users can perform standard DR-SIP docking protocols with the CLI while more advanced users can import the specific modules and functions to perform their own customized docking protocol.

The source code and documentation on how to use the DR-SIP package are provided at https://github.com/capslockwizard/drsip. The packages are also distributed through the PyPi and Anaconda Cloud repositories.

## RESULTS

### Statistical Analysis of 115 Structurally Solved α-Helical HoTPs

To observe how common C_n_ symmetric structures are and how well the properties of C_n_ symmetry holds for α-helical HoTPs, 152 non-redundant structures containing HoTPs were taken from the mpstruct database (http://blanco.biomol.uci.edu/mpstruc/), see Supplementary Table file. Out of the 152 structures, 23 of them are monomers or do not contain a transmembrane domain. The remaining 129 structures (64% are observed within asymmetric units) contain 118 structures (includes two dihedral symmetric structures, see Supplementary Methods) with parallel orientation (N-termini of constituent monomers face the same side of the membrane) and 11 anti-parallel structures (see Supplementary Results). The 118 parallel structures which constitute 91.5% of the 129 HoTPs were used for subsequent statistical analyses. The anti-parallel structures are not included because most have not been verified as their functional forms through supporting experiments (see Supplementary Results).

The distribution of the sizes of the 118 parallel HoTPs (Fig. 3A) show that most are dimers (47 HoTPs), trimers (28 HoTPs) and tetramers (26 HoTPs), for a total of 101 HoTPs (85.6%) out of the 118 HoTPs.

We further identified 115 of the 118 parallel HoTPs to be C_n_ symmetric (Fig. 3B-E, C_n_ symmetry RMSD <2Å) concluding that most of the HoTPs (89.1% or 115/129) are C_n_ symmetric and parallel in orientation.

#### Decomposition of the Deviation from Ideal C_n_ Symmetry

There are two potential contributors to the deviation from ideal C_n_ symmetry. The first is the conformational difference between monomers in the complex. The other is the translational and rotational differences between a docking pose and its closest ideal C_n_ symmetric pose.

Our results show that conformational differences between monomers is not a factor preventing the formation of C_n_ symmetry. All the monomers in the 118 parallel HoTPs have small structural differences (Fig. 3B) – most (~90%) are less than 1Å, while all are less than 3Å.

On the other hand, the C_n_ symmetry RMSD (Fig. 3C) measures the translational and rotational differences between the ideal C_n_ symmetric pose and the pose from the X-ray structure for all unique pairs of neighboring monomers. When the C_n_ symmetry RMSD is <2.0Å (Fig. 3C, red line and inset), the pose is classified as C_n_ symmetric. This is the same cutoff used in the SIP filter. Among the 118 parallel HoTPs, 115 (97.5%) were classified as C_n_ symmetric.

The cutoff of <2.0Å was chosen because it is the midpoint between 1.6Å and 2.4Å where there are no structures (Fig. 3C). There are only three HoTPs (Supplementary Fig. S4 and Table S5) that have C_n_ symmetry RMSD >2.4Å and upon further examination were found to be non-Cn symmetric (see Supplementary Results for details on the three HoTPs). Figure 4 compares the X-ray structures of the two HoTPs closest to the 2.0Å cutoff, one above it and the other below it, with their respective ideal C_n_ symmetric complex. The ExbB/ExbD complex (Fig. 4A) which is below the cutoff does not deviate much (see next section) from the ideal C_n_ symmetric complex while the multidrug efflux pump subunit AcrB (Fig. 4B) shows a visibly larger deviation (see next section).

#### Decomposition of Deviations from Ideal C_n_ Symmetry into Translation and Rotational Differences

Deviations from ideal C_n_ symmetry are decomposed into translational (shift along AOS, sAOS) and rotational (deviation from ideal rotation angle, devRot) differences. The statistics show that all 115 C_n_ symmetric HoTPs have <11° deviation from the ideal rotation angle (94% is <2°, Fig. 1D) and <0.7Å shift along AOS (95% is <0.3Å, Fig. 1E).

All the three non-C_n_ symmetric structures (Supplementary Table S5) have ≥0.7Å sAOS while two out of the three (acid-sensing ion channel and the trimeric AcrB [Fig. 4B]) have ≥11° devRot with the remaining one >6.0°. For comparison, the ExbB/ExbD complex (Fig. 4A) has 0.5Å sAOS and 4.0° devRot which are smaller than the three non-C_n_ symmetric structures.

#### Tilt Angles Distribution

The distribution of tilt angles (Fig. 1F) show that all the 115 structures have tilt angles <50°, while 90% of them is <30°; hence 30° (red line in Fig. 1F) is used as the tilt angle cutoff for the SIP filter.

### SIP Filter Performance Without Distance Restraints

Experimentally measured distances of interacting monomers in a HoTP complex are not commonly available. Even without distance restraints (DR), SIP alone can predict a near-native pose for every ~3/4 HoTPs (73%, Table 1).

**Table 1.** Percentage of 115 HoTPs Whose Near-Native Pose Can Be Recovered by SIP or ZDOCK Without Distance Restraints.

|  | **Top-20 (%)** | **Top-10 (%)** | **Top-5 (%)** | **Top-3 (%)** |
| --- | --- | --- | --- | --- |
| **ZDOCK<sup>a,c</sup>** | 47.8 | 45.2 | 40.9 | 39.1 |
| **DR-SIP (Ex. DR)<sup>b,c</sup>** | 73.0 | 64.3 | 52.2 | 42.6 |
| **ZDOCK Rank (Ex. Dimers)<sup>d</sup>** | 32.4 | 27.9 | 23.5 | 23.5 |
| **DR-SIP (Ex. DR) Rank (Ex. Dimers)<sup>d</sup>** | 67.6 | 57.4 | 48.5 | 39.7 |

<sup>a</sup> For each HoTP, the original ZDOCK rank of the unfiltered 54,000 docking poses.
<sup>b</sup> For each HoTP, the SIP filter ( $C_n$ symmetry $>2\text{\AA}$ , tilt angle $>35^\circ$ ) and clustering are applied on the 54,000 docking poses. The clusters and their respective representative poses are ranked by size in descending order. See **Methods** and **Fig. 2**.
<sup>c</sup> Percentage of the 115 HoTPs.
<sup>d</sup> Percentage of the remaining 68 HoTPs after excluding all 47 dimers.

The 115 HoTP poses from the dataset were docked with ZDOCK and filtered by SIP. SIP increases the percentage of HoTPs that can be found within the top-3, 5, 10 and 20 ranking results (Table 1). The largest increase comes from improving the number of HoTPs with near-native pose(s) within the top-20 (DR-SIP: 73% of 115 HoTPs, ZDOCK: 47.8%) and top-10 (DR-SIP: 64.3%, ZDOCK: 45.2%) results.

Since the dataset is dominated by dimers (40.9% of HoTP, Fig. 3A), it is possible that SIP is just good at predicting dimers. By excluding dimers from the statistics, we show that SIP performs just as well (top-20: 67.6%, top-10: 57.4%) for higher order oligomers, while the performance of ZDOCK drops to 32 and 28%, respectively (Table 1).

Near-native docking poses are identified by comparing each docking pose to their respective reference X-ray structure by a modified CAPRI criteria (Méndez *et al*., 2003) for HoTPs which assigns each pose into one of four ranks: High, Medium, Acceptable and Incorrect, respectively (see Supplementary Methods and Table S6). Docking poses that are not “Incorrect” ([<10% native contacts] or [>4.0 iRMSD and >10.0 lRMSD] or [have a different order of symmetry with the reference structure]) are considered as near-native poses.

### Performance of DR-SIP with Distance Restraints: Near-Native Poses Within the Top-2 Ranking Poses

The performance of DR-SIP when distance restraints are available is evaluated by applying the docking protocols (Fig. 2) to one soluble protein system (Syt1-SNARE) and three HoTP systems (MscL, *Vr*H^+^-PPase and *Ct*H^+^-PPase). The results are compared to DR-SIP without distance restraints and the original ZDOCK ranking (Table 2).

**Table 2.** Improvement of the Ranking of the Top-Ranked Near-Native Docking Poses with DR-SIP.

|  | Syt1-SNARE | MscL | CtH<sup>+</sup>-PPase | VrH<sup>+</sup>-PPase |
| --- | --- | --- | --- | --- |
| **ZDOCK<sup>a</sup>** | 62 | 2231 | 1 | 1 |
| **SIP only (Ex. DR)<sup>b</sup>** | - | 7 | 1 | 1 |
| **DR-SIP<sup>c</sup>** | 1 | 2 | 1 | 1 |

Shows the top ranking near-native pose for four systems: Syt1-SNARE, MscL, CtH<sup>+</sup>-PPase and VrH<sup>+</sup>-PPase.
<sup>a</sup> The original ZDOCK rank of the 54,000 unfiltered poses for all four systems.
<sup>b</sup> DR-SIP ranking obtained by applying only the SIP filter ( $C_n$ symmetry $>2\text{\AA}$ , tilt angle $>35^\circ$ ), just like in **Table 1**.
<sup>c</sup> DR-SIP ranking obtained by applying both the DR filter ( $\leq 0.3$ correlation) and SIP filter ( $C_n$ symmetry $>2\text{\AA}$ , tilt angle $>35^\circ$ ), see **Methods** and **Fig. 2**.

A near-native pose of *Vr*H^+^-PPase and *Ct*H^+^-PPase can be found within the top-20 ZDOCK ranked poses but not for Syt1-SNARE and MscL. Consistent with the results of the previous section, applying SIP without the DR filter on MscL increases the chance to find a near-native pose within the top-10. Indeed, one is found at the 7^th^-ranked pose. The ranking is further improved to 2^nd^ (Fig. 5A,B) when the DR filter is added. While for the soluble system, Syt1-SNARE (a hetero-oligomer where C_n_ symmetry does not apply), applying only the DR filter, the top-ranked pose (Fig. 5B) is the near-native pose.

**Fig. 5.**
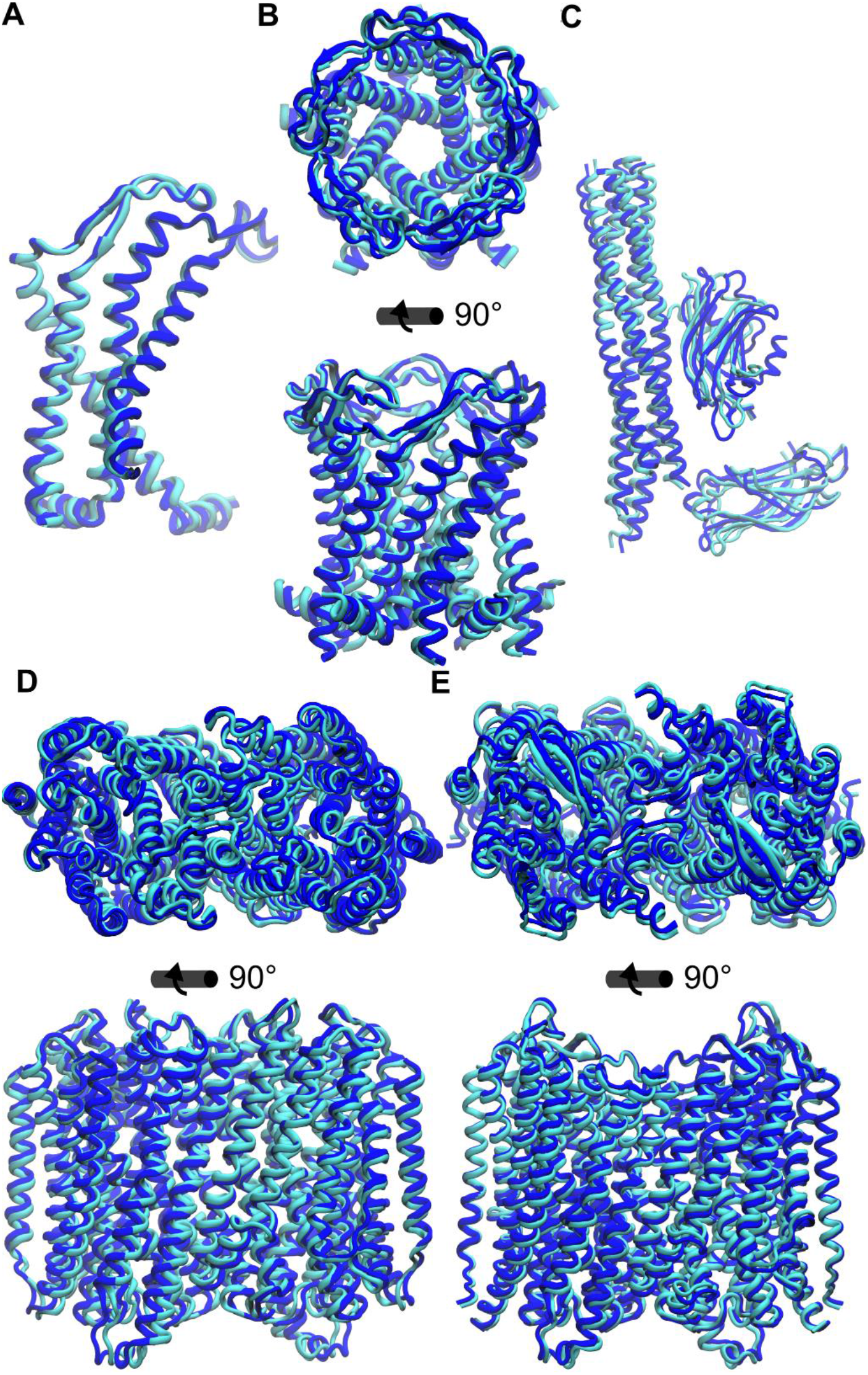
The top-ranked near-native docking pose (cyan) predicted by DR-SIP superimposed onto the reference structure (blue) for all four systems with distance restraints. **(A-B)** MscL: 2^nd^ ranked pose with the reference structure from PDB ID: 2OAR. (**A**) The docking pose (dimer) superimposed onto chain A and B of the reference structure. (**B**) The predicted pentamer complex with the reference structure. (**C**) Syt1-SNARE complex: the 1^st^ ranked DR-SIP pose with the reference structure PDB ID: 5CCG. (**D**) *Ct*H^+^-PPase: 1^st^ ranked pose with the homology modeled structure based on the template PDB ID: 4A01. (**E**) *Vr*H^+^-PPase: 1^st^ ranked pose with the reference structure PDB ID: 4A01.

DR-SIP performs equivalently for systems that are ranked 1^st^ by ZDOCK such as *Vr*H^+^-PPase and *Ct*H^+^-PPase. We want to emphasize that DR-SIP does not rely on the ranking of ZDOCK, or the ranking of any other software that generates the initial pool of docking poses. For “tougher” systems, DR-SIP enriches the near-native poses such that they can be found within the top-2 results (Table 2 and Fig. 5).

### Contribution of the Filters and Clustering to the Enrichment of the Near-Native Poses

Among the 54,000 docking poses generated by ZDOCK for each of the four systems, <0.3% (Table 3) of the poses are near-native poses. The docking protocols enriched the near-native poses up to 1250-fold such that at least one can be found within the top-2 ranking results (Table 2 and 3).

**Table 3.** Contribution of the DR-SIP Filters to Enriching the Near-Native Poses Generated by ZDOCK.

|  | % Native<sup>c</sup> | Enrichment<sup>d</sup> | Native Poses |  |  | Incorrect | Total |
| --- | --- | --- | --- | --- | --- | --- | --- |
|  |  |  | High | Medium | Acceptable |  |  |
| Syt1-SNARE |  |  |  |  |  |  |  |
| Unfiltered | 0.25 | 1.00 | 3 | 23 | 108 | 53866 | 54000 |
| All Filters<sup>a</sup> | 2.70 | 10.89 | 3 | 23 | 108 | 4824 | 4958 |
| Filtered<sup>a</sup> & Clustered | 1.65 | 6.64 | 0 | 1 | 6 | 418 | 425 |
| Top 2 with DR | 50.00 | 201.49 | 0 | 0 | 1 | 1 | 2 |
| MscL |  |  |  |  |  |  |  |
| Unfiltered | 0.04 | 1.00 | 4 | 11 | 5 | 53980 | 54000 |
| All Filters<sup>b</sup> | 9.73 | 243.25 | 4 | 7 | 0 | 102 | 113 |
| Ex. C<sub>n</sub> Symm. RMSD | 0.56 | 14.00 | 4 | 8 | 1 | 2294 | 2307 |
| Ex. Tilt Angle | 1.38 | 34.50 | 4 | 8 | 1 | 931 | 944 |
| Ex. DR Filter | 2.35 | 58.75 | 4 | 7 | 0 | 457 | 468 |
| Ex. DR Filter & Clustered | 4.17 | 104.25 | 0 | 1 | 0 | 23 | 24 |
| Filtered<sup>b</sup> & Clustered | 11.11 | 277.75 | 0 | 1 | 0 | 8 | 9 |
| Top 7 (Ex. DR) | 14.29 | 357.25 | 0 | 1 | 0 | 6 | 7 |
| Top 2 (Ex. DR) | 0.00 | 0.00 | 0 | 0 | 0 | 2 | 2 |
| Top 7 with DR | 14.29 | 357.25 | 0 | 1 | 0 | 6 | 7 |
| Top 2 with DR | 50.00 | 1250.00 | 0 | 1 | 0 | 1 | 2 |
| CtH<sup>+</sup>-PPase |  |  |  |  |  |  |  |
| Unfiltered | 0.14 | 1.00 | 0 | 29 | 45 | 53926 | 54000 |
| All Filters<sup>b</sup> | 26.15 | 186.79 | 0 | 19 | 15 | 96 | 130 |
| Ex. C<sub>n</sub> Symm. RMSD | 2.12 | 15.14 | 0 | 29 | 45 | 3417 | 3491 |
| Ex. Tilt Angle | 9.69 | 69.21 | 0 | 19 | 15 | 317 | 351 |
| Ex. DR Filter | 10.56 | 75.43 | 0 | 19 | 15 | 288 | 322 |
| Ex. DR Filter & Clustered | 3.57 | 25.50 | 0 | 1 | 0 | 27 | 28 |
| Filtered<sup>b</sup> & Clustered | 12.50 | 89.29 | 0 | 1 | 0 | 7 | 8 |
| Top 7 (Ex. DR) | 14.29 | 102.07 | 0 | 1 | 0 | 6 | 7 |
| Top 2 (Ex. DR) | 50.00 | 357.14 | 0 | 1 | 0 | 1 | 2 |
| Top 7 with DR | 14.29 | 102.07 | 0 | 1 | 0 | 6 | 7 |
| Top 2 with DR | 50.00 | 357.14 | 0 | 1 | 0 | 1 | 2 |
| VrH<sup>+</sup>-PPase |  |  |  |  |  |  |  |
| Unfiltered | 0.17 | 1.00 | 4 | 29 | 58 | 53909 | 54000 |
| All Filters<sup>b</sup> | 16.67 | 98.06 | 3 | 13 | 15 | 155 | 186 |
| Ex. C<sub>n</sub> Symm. RMSD | 2.00 | 11.76 | 4 | 29 | 58 | 4451 | 4542 |
| Ex. Tilt Angle | 7.49 | 44.06 | 3 | 13 | 15 | 383 | 414 |
| Ex. DR Filter | 7.60 | 44.71 | 3 | 13 | 15 | 377 | 408 |
| Ex. DR Filter & Clustered | 2.33 | 13.71 | 0 | 1 | 0 | 42 | 43 |
| Filtered<sup>b</sup> & Clustered | 6.25 | 36.76 | 0 | 1 | 0 | 15 | 16 |
| Top 7 (Ex. DR) | 14.29 | 84.06 | 0 | 1 | 0 | 6 | 7 |
| Top 2 (Ex. DR) | 50.00 | 294.12 | 0 | 1 | 0 | 1 | 2 |
| Top 7 with DR | 14.29 | 84.06 | 0 | 1 | 0 | 6 | 7 |
| Top 2 with DR | 50.00 | 294.12 | 0 | 1 | 0 | 1 | 2 |

<sup>a</sup> Using only the DR filter ( $\leq 0.3$ correlation).
<sup>b</sup> Using both the DR filter ( $\leq 0.3$ correlation) and SIP filter ( $C_n$ symmetry $> 2\text{\AA}$ , tilt angle $> 35^\circ$ ).
<sup>c</sup> % Native = (High + Medium + Acceptable) Poses / Total Poses \* 100.
<sup>d</sup> Enrichment = (% Native) / (% Native when unfiltered). For example, MscL's "Top 2 with DR" enrichment of 1250.0 is obtained by dividing the 50% near-native poses with the ~0.04% near-native poses in the unfiltered data.

The contribution of the filters is measured by excluding one of the four filters/criteria and comparing the decrease in the enrichment of near-native poses relative to the enrichment when using all filters/criteria (Table 3). The larger the drop in the enrichment, the larger the contribution of that filter.

The C_n_ symmetry criterion has the largest contribution to the enrichment resulting in a drop of ~90% (in Table 3, [**Ex. C_n_ Symm. RMSD/All Filters - 1**] x 100%) when excluded. On the other hand, the tilt angle criterion and relative distance filters are of equal importance, reducing the enrichment by ~55-85% when excluded.

After applying all the filters, the near-native poses are significantly enriched (up to 243-fold) and make up 10%-25% of the remaining poses. The near-native poses are further enriched by clustering of the poses (see METHODS) such that the near-native pose can be found within the top-2 results.

A near-native pose of MscL cannot be found in the top-2 results without the DR filter. MscL starts off with 3 to 4-fold fewer near-native poses (0.04% out of 54,000 poses) compared to the other two HoTPs (~0.16%). Without the DR filter, the SIP filter can only enrich the near-native poses to 2.35% of poses. This is enough to obtain a near-native pose at the 7^th^-ranked pose (Table 2). A 4-fold enrichment by the DR filter pushes the final rank to 2^nd^ (Table 2).

For the soluble case, the Syt1-SNARE complex, the DR filter enriches the near-native poses by ~11-fold. Clustering and ranking the remaining poses results in a 1^st^-ranked near-native pose.

### Performance of DR-SIP with GRAMM-X

To examine how DR-SIP performs with other docking software, DR-SIP is applied to the docking poses generated from GRAMM-X for all four systems. From the maximum 300 poses that could be obtained for each system, GRAMM-X results contain the near-native pose(s) for MscL, *Vr*H^+^-PPase and *Ct*H^+^-PPase but not for the Syt1-SNARE complex. Applying DR-SIP (excluding Syt1-SNARE) enriches the near-native poses to ~33-50% of remaining poses from the unfiltered 0.3-2.7% (Table 4), performing similarly to the ZDOCK results (Table 3), which shows the generality of DR-SIP’s performance as different docking packages are used.

**Table 4.** GRAMM-X Docking Results Filtered by the DR-SIP Protocols.

| System | % Native Poses<br>[Before Filter] | % Native Poses<br>[After Filter]<sup>a</sup> | Enrichment<sup>b</sup> |
| --- | --- | --- | --- |
| **MscL** | 2.7 (8/300) | 33.3 (2/6) | 12.3 |
| **Ct-PPase** | 0.3 (1/300) | 50.0 (1/2) | 166.7 |
| **Vr-PPase** | 0.3 (1/300) | 33.3 (1/3) | 111.0 |
| **Syt1-SNARE** | 0.0 (0/300) | 0.0 (0/300) | - |

<sup>a</sup> Using both the DR filter ( $\leq 0.3$ correlation) and SIP filter ( $C_n$ symmetry $> 2\text{\AA}$ , tilt angle $> 35^\circ$ ). No clustering was performed because there are too few poses after filtering.
<sup>b</sup> Enrichment = % Native Poses [After Filter] / % Native Poses [Before Filter].

## DISCUSSIONS

The distribution of the “*n*” of C_n_ symmetric HoTPs (Fig. S5A and Table S7) is not uniformly distributed, with dimers and trimers dominating the population. Such a distribution could occur from a natural outcome of chemical/geometric complementarity of the constituent monomers (physical reason) or meeting the functional needs of life (biological reason).

To see whether a purely physical model can reproduce the observed distribution of “*n*”, we perform molecular docking with ZDOCK on three HoTP systems (MscL, *Vr*H^+^-PPase and *Ct*H^+^-PPase) and use the C_n_ symmetry criterion to select only symmetric poses. ZDOCK, as an off-lattice model (Huang, 2014), uniformly samples the translational and rotational degrees of freedom and returns the top-54,000 poses with the best interaction scores between two identical monomers.

Our results show that the proportion of C_2_’s is over-represented, ~73.8% of total, (Fig. S5A, S5C and Table S7) relative to the observed 40.9% in the HoTP dataset. Not unique to HoTPs, C_n_ symmetric *soluble proteins* (Pentameric Capsid Protein [PDB ID: 4DMI] and GH3 Beta-Glucosidase [PDB ID: 5XXL]), selected for having a similar size as MscL and *Vr*H^+^-PPase/*Ct*H^+^-PPase, respectively (see **Supporting Methods** for details), also show C_2_’s are overpopulated (~69.5%).

The discrepancy between our results and the observed distribution could be due the lack of consideration given to the constraints imposed on HoTPs as they span the membrane bilayer. One such constraint is the need for transmembrane regions of the monomers in HoTPs to be embedded in the membrane. When the transmembrane regions are longer than the thickness of the membrane, the monomers must undergo a tilt to fully embed these regions (Park and Opella, 2005; Kim and Im, 2010; de Jesus and Allen, 2013). Our statistics (Fig. 3F) show that a relatively small tilt angle (<35°; compared to the mean angle of 57.3° (Li *et al*., 2014) for two random lines in space) is preferred, which implies that most of the transmembrane regions are not too long. While, large tilt angles imply that the transmembrane region is longer, and this might be disadvantageous to the formation of sufficient binding interface in C_2_ complexes and/or cost-effectiveness of protein synthesis.

After applying the tilt angle criterion (<35°) to the C_n_ symmetric ZDOCK poses, we observe a distribution that is similar (C2 make up ~53.6%) to the HoTP dataset (Fig. S5B, S5D and Table S7). Therefore, the tilt angle constraint is one of the major determinants of the observed distribution. However, there is no straightforward physical explanation of why tilt angles are predominantly populated over 7° and 20° (Fig. 3F). Therefore, we surmise that this “selection” is made for functional reasons such as proper functional dynamics or genetic reasons to provide evolutionary advantages.

The monomers in a C_2_ complex share the *same binding interface* but the monomers in the other C_n_’s interact at *different interfaces*. Therefore, any interaction between residues at a C_2_ interface comes in pairs (Monod *et al*., 1965). The interaction energy of good/bad interactions are doubled resulting in a larger variance of the interaction energies for C2’s (André *et al*., 2008). Thus, we’re are more likely to find C2’s with very favorable interaction energies compared to other types of dimers (André *et al*., 2008). Since molecular docking returns poses with the most favorable interactions, we expect and indeed observe that C2’s are enriched compared to other Cn’s (Fig. S5A and Table S7). Thus, physical interactions can explain why C_2_ HoTPs are so common. However, only a subset of these C_2_’s within the range of allowable tilt angles were chosen by evolution for functional purposes.

Taken together, DR-SIP incorporates geometric restraints such as C_n_ symmetry and tilt angle for HoTPs while leveraging experimentally determined distance restraints providing a reliable opportunity to computationally assemble near-native quaternary structures of hetero-/homo-oligomers.

## Acknowledgments

Vast computational resources for this project are provided by National Center for High-performance Computing (NCHC) of National Applied Research Laboratories (NARLabs), Taiwan. J.C. acknowledges financial support from Taiwan International Graduate Program (TIGP), Academia Sinica, Taiwan, as well as θr. Tzyy-Jen Chiou’s support (AS 103-TP-B11). This work is supported by the Ministry of Science and Technology, Taiwan (103-2627-M-007-001 and 104- 2113-M-007-019 to L.-W. Y.).

