## Supplementary Data for "DR-SIP: Protocols for Higher Order Structure Modeling with Distance Restraints- and Cyclic Symmetry-Imposed Packing"

### Supplementary Figures

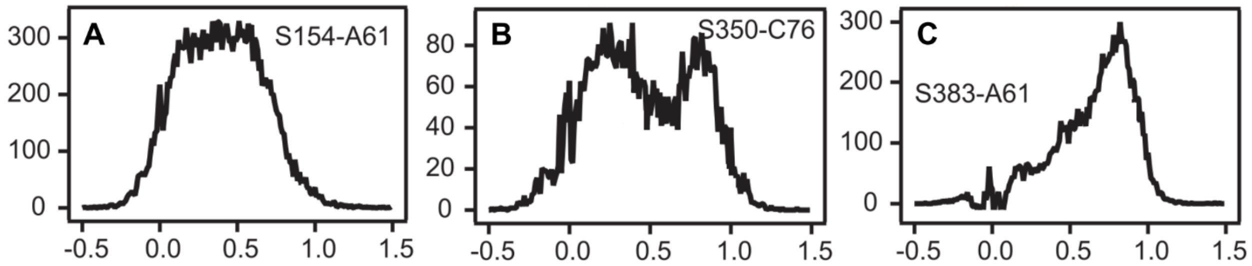

**Fig. S1.** Examples of smFRET transfer efficiency histograms that do not contain a single Gaussian-like peak. (**A**) Broad peaks, where the difference between the maximum and minimum transfer efficiency of the peak is >50%. (**B**) Multiple peaks. (**C**) Non-Gaussian distribution. Any smFRET pairs between Syt1 and SNARE complex whose transfer efficiency histogram contains one or more of the above properties were removed (**Table S1**). The smFRET transfer efficiency histograms were taken from the Supplementary Data of Zhou *et al.* (Zhou *et al.*, 2015) with modifications.

```
MscL-Ecoli            1  MSIIKEFREFAMRGNVVDLAVGVIIGAAFGKIVSSLVADIIMPPLGLL..IGGIDFKQFA
MscL-Mtuberculosis    1  ..MLKGFKEFLARGNIVDLAVAVVIGTAFTALVTKFTDSIITPLINRIGVNAQSDVGILR
MscL-Mleprae          1  ..MFRGFKEFLSRGNIVDLAVAVVIGTAFTALITKFTDSIITPLINRVGVNQQTNISPLR
MscL-Bsubtilis        1  ..MWNEFKAFAMRGNIVDLAIGVVIGGAFGKIVTSLVNDIIMPLVGLL..LGGLDFSGLS
MscL-Hinfluenzae      1  MNFIKEFREFAMRGNVVDMAVGVIIGSAFGKIVSSLVSDIFMPVLGIL..TGGIDFKDMK

MscL-Ecoli           59  VTLRDAQGDIPAVVMHYGVFIQNVFDFLIVAFAIFMAIKLINKLNRKKEEPAAAP..APT
MscL-Mtuberculosis   59  IGIGGGQ......TIDLNVLLSAAINFFLIAFAVYFLVVLPYNTLRKKGEVEQPG....D
MscL-Mleprae         59  IDIGGDQ......AIDLNIVLSAAINFLLIALVVYFLVVLPYTTIRKHGEVEQFDTDLIG
MscL-Bsubtilis       57  FTFGDA.......VVKYGSFIQTIVNFLIISFSIFIVIRTLNGLRRKKEAEEEAAEEAVD
MscL-Hinfluenzae     59  FVLAQAQGDVPAVTLNYGLFIQNVIDFIIIAFAIFMMIKVINKVRKPEEKKPVPK.....

MscL-Ecoli          117  KEEVLLTEIRDLLKEQNNRS.........................
MscL-Mtuberculosis  109  TQVVLLTEIRDLLAQTNGDSPGRHGGRG..TPSPTDGPRASTESQ
MscL-Mleprae        113  NQVVLLAEIRDLLAQSNGAPSGRHVDTADLTPTPNHEPRADT...
MscL-Bsubtilis      110  AQEELLKEIRDLLKQQAKSPE........................
MscL-Hinfluenzae    114  .AETLLTEIRDLLKNK.............................
```

**Fig. S2.** Mapping (boxes) of the residues of *E. coli* MscL measured in the smFRET experiment by Wang *et al.* (Wang *et al.*, 2014) to the *M. tuberculosis* MscL based on the multiple sequence alignment of five MscL proteins (Edgar, 2004). The sequence identity between the two sequences is 32.6%. See **Supplementary Methods** and **Table S3** for the details.

```
Ct-PPase    1  MESFIVYSVLAGVIALIFAFMLSSFISKE.......................SAG....N
Vr-PPase   12  ....EILIPVCAVIGIAFALFQWLLVSKVKLSAVRDASPNAAAKNGYNDYLIEEEEGIND

Ct-PPase   34  ....ERMQEIAGHIHDGAMAFLKTEYKYLTGFIVIVTVILAIFVGW..............
Vr-PPase   68  HNVVVKCAEIQNAISEGATSFLFTEYKYVGIFMVAFAILIFLFLGSVEGFSTSPQACSYD

Ct-PPase   76  .............QTAACFILGAIFSIFAGYFGMNVATKANVRTAEAARHSQGKALNIAF
Vr-PPase  128  KTKTCKPALATAIFSTVSFLLGGVTSLVSGFLGMKIATYANARTTLEARKGVGKAFITAF

Ct-PPase  123  SGGAVMGMSVVGLGVVGIGIMYYIFG.....G...NMEFVTGFGLGASSIALFARVGGGI
Vr-PPase  188  RSGAVMGFLLAANGLLVLYIAINLFKIYYGDDWGGLFEAITGYGLGGSSMALFGRVGGGI

Ct-PPase  175  YTKAADVGADLVGKVEAGIPEDDPRNPAVIADNVGDNVGDVAGMGADLFESYVGSIISAL
Vr-PPase  248  YTKAADVGADLVGKVERNIPEDDPRNPAVIADNVGDNVGDIAGMGSDLFGSYAESSCAAL

Ct-PPase  235  TLGTVVYAN....KEGVMFPLILSSIGIVASIIGILFS....RKSKAKDPQKALNTGTYI
Vr-PPase  308  VVASISSFGLNHELTAMLYPLIVSSVGILVCLLTTLFATDFFEIKAVKEIEPALKKQLVI

Ct-PPase  287  GGIIVIVSAAILSNTIFG...............NLKAFFAVASGLVVGMIIGKITEMYTS
Vr-PPase  368  STVLMTIGVAVVSFVALPTSFTIFNFGVQKDVKSWQLFLCVAVGLWAGLIIGFVTEYYTS

Ct-PPase  332  DAYSSVQKIANQSETGPATTIISGLAVGMYSTLWPIVLISIGVLVSFFVMGGGSNAMVGL
Vr-PPase  428  NAYSPVQDVADSCRTGAATNVIFGLALGYKSVIIPIFAIAISIFVSFTFA........AM

Ct-PPase  392  YGISLAAVGMLSTTGLTVAVDAYGPIADNAGGIAEMSELPHEVREITDKLDSVGNTTAAI
Vr-PPase  480  YGIAVAALGMLSTIATGLAIDAYGPISDNAGGIAEMAGMSHRIRERTDALDAAGNTTAAI

Ct-PPase  452  GKGFAIGSAALTALSLFASYAQATELESIDILNTVTLVGLFIGAMLPFLFGALTMESVGK
Vr-PPase  540  GKGFAIGSAALVSLALFGAFVSRASITTVDVLTPKVFIGLIVGAMLPYWFSAMTMKSVGS

Ct-PPase  512  AANEMIEEVRRQFKTIPGIMEGKATPDYKKCVDISTAAAIREMILPGVLAIVVPVAMGLL
Vr-PPase  600  AALKMVEEVRRQFNTIPGLMEGTAKPDYATCVKISTDASIKEMIPPGALVMLTPLVVGIL

Ct-PPase  572  LGKEALGGLLAGALVSGVLVGILMSNAGGAWDNAKKYIEGGAH......GGKGSEAHKAA
Vr-PPase  660  FGVETLSGVLAGSLVSGVQIAISASNTGGAWDNAKKYIEAGASEHARSLGPKGSDCHKAA

Ct-PPase  626  VVGDTVGDPFKDTSGPSMNILIKLMTIVSLVFAPVVLQYGGILLNLIK
Vr-PPase  720  VIGDTIGDPLKDTSGPSLNILIKLMAVESLVFAPFFATHGGLLFK...
```

**Fig. S3.** Mapping (boxes) of the residues in *Ct*H$^{+}$-PPase measured in the smFRET experiment (Huang *et al.*, 2013) to the corresponding *Vr*H$^{+}$-PPase residues based on the pairwise sequence alignment (45.1% identity) generated by SwissModel (Biasini *et al.*, 2014) for homology modeling. See **Supplementary Methods** and **Table S4** for details.

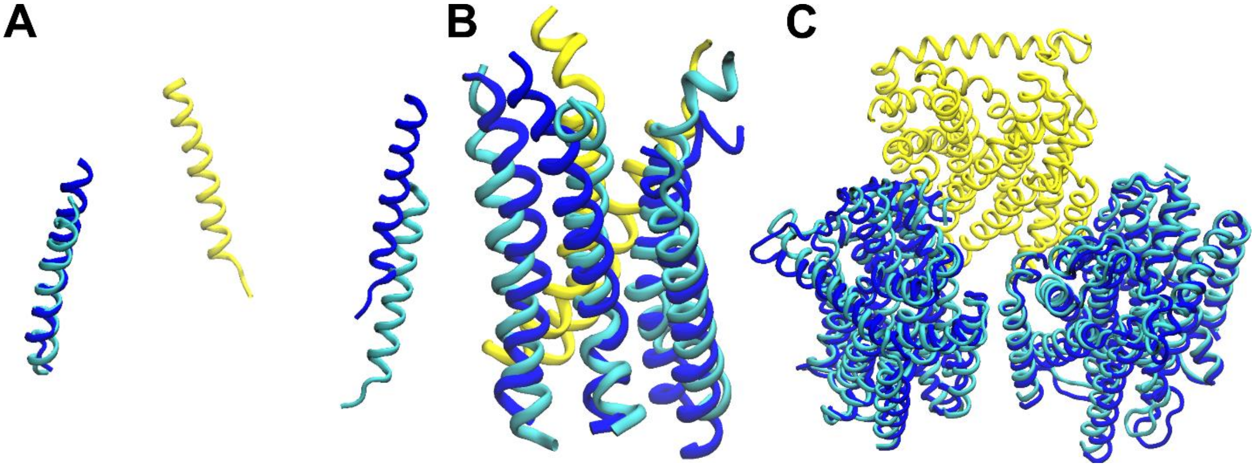

**Fig. S4.** X-ray structures (blue) of the three non-$C_n$ symmetric trimeric complexes superimposed onto their ideal $C_3$ symmetric complexes (cyan) at a single monomer (in yellow). (**A**) Human σ1 receptor protein (PDB ID: 5HK1) with the $C_n$ symmetry RMSD of 21.1Å. (**B**) Acid-sensing ion channel (PDB ID: 2QTS), RMSD: 4.6Å. (**C**) Multidrug efflux pump subunit AcrB (PDB ID: 2GIF), RMSD: 2.5Å. See details in **Supplementary Table S5**.

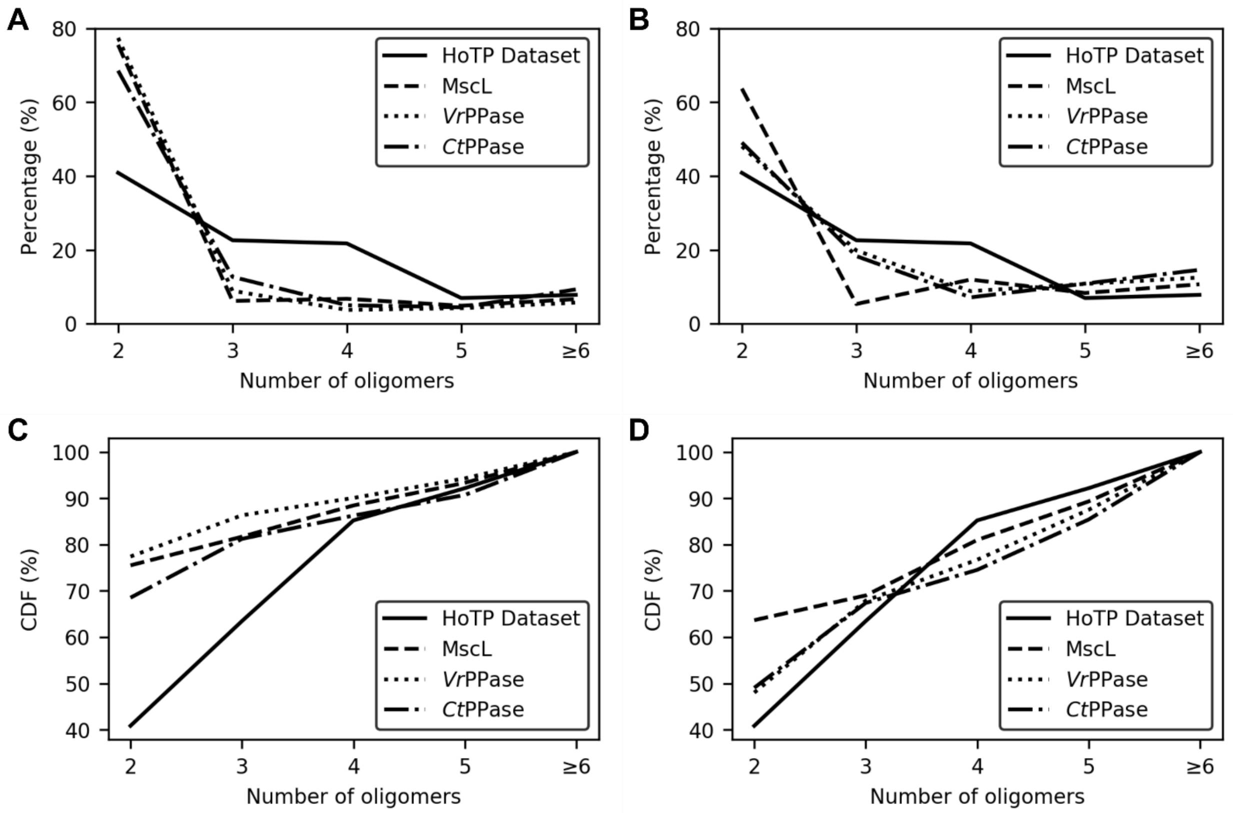

**Fig. S5**. The size distribution (**A and B**, counts normalized to percentage of total poses) and cumulative distribution function (CDF) (**C and D**) of the $C_n$ symmetric HoTP dataset (X-ray-solved structures) and the docking poses from the MscL, *Vr*$H^+$-PPase and *Ct*$H^+$-PPase systems. (**A and C**) Filtered with only the $C_n$ symmetry criterion ($C_n$ RMSD > 2Å). (**B and D**) Filtered with the $C_n$ symmetry and tilt angle (tilt angle > 35°) criteria.

### Supplementary Tables

#### Table S1. smFRET data for the Syt1-SNARE complex.

| Monomer 1 | Resid 1 | Monomer 2 | Resid 2 | Include | Reason for Exclusion[a] |
|---|---|---|---|---|---|
| S25 | 20 | Syt1 | 140 | Yes | - |
| S25 | 20 | Syt1 | 154 | Yes | - |
| S25 | 20 | Syt1 | 368 | Yes | - |
| S25 | 20 | Syt1 | 383 | Yes | - |
| S25 | 76 | Syt1 | 140 | Yes | - |
| S25 | 20 | Syt1 | 269 | No | Multiple peaks |
| S25 | 20 | Syt1 | 350 | No | Peak cover >50 % |
| S25 | 76 | Syt1 | 154 | No | Multiple peaks |
| S25 | 76 | Syt1 | 269 | No | Multiple peaks |
| S25 | 76 | Syt1 | 350 | No | Multiple peaks |
| S25 | 76 | Syt1 | 368 | No | Multiple peaks |
| S25 | 76 | Syt1 | 383 | No | Multiple peaks |
| S25 | 139 | Syt1 | 140 | No | Peak cover >50 % |
| S25 | 139 | Syt1 | 154 | No | Multiple peaks |
| S25 | 139 | Syt1 | 269 | No | Multiple peaks |
| S25 | 139 | Syt1 | 350 | No | Non-Gaussian |
| S25 | 139 | Syt1 | 368 | No | Non-Gaussian |
| S25 | 139 | Syt1 | 383 | No | Multiple peaks |
| S25 | 197 | Syt1 | 140 | No | Non-Gaussian |
| S25 | 197 | Syt1 | 154 | No | Peak cover >50 % |
| S25 | 197 | Syt1 | 269 | No | Peak cover >50 % |
| S25 | 197 | Syt1 | 350 | No | Non-Gaussian |
| S25 | 197 | Syt1 | 368 | No | Multiple peaks |
| S25 | 197 | Syt1 | 383 | No | Multiple peaks |
| Sb | 61 | Syt1 | 140 | No | Non-Gaussian |
| Sb | 61 | Syt1 | 154 | No | Peak cover >50 % |
| Sb | 61 | Syt1 | 269 | No | Non-Gaussian |
| Sb | 61 | Syt1 | 350 | No | Multiple peaks |
| Sb | 61 | Syt1 | 368 | No | Non-Gaussian |
| Sb | 61 | Syt1 | 383 | No | Non-Gaussian |

[a] smFRET residue pairs that have only a single peak in their transfer efficiency histograms were chosen. Histograms showing multiple peaks, non-Gaussian distribution or peaks that are broad were excluded (see details and examples in **Supplementary Fig. S1**). The smFRET data are from Choi *et al.* (Choi *et al.*, 2010).

#### Table S2. Syt1-SNARE complex's smFRET data.

| Monomer 1 | Resid 1 | Monomer 2 | Resid 2 | smFRET Dist. (nm)[a,c] | Ref Structure Dist. (nm)[b,c] | DR-SIP Pred. Pose Dist. (nm)[d] |
|---|---|---|---|---|---|---|
| S25 | 20 | Syt1 | 140 | 6.4 | 4.5 | 4.3 |
| S25 | 20 | Syt1 | 154 | 5.9 | 2.3 | 3.1 |
| S25 | 20 | Syt1 | 368 | 6.7 | 5.0 | 5.7 |
| S25 | 20 | Syt1 | 383 | 6.2 | 5.0 | 4.8 |
| S25 | 76 | Syt1 | 140 | 7.1 | 8.7 | 8.8 |

[a] smFRET-measured distances are from Choi *et al.* (Choi *et al.*, 2010).
[b] The distances are between Cα atoms. The reference structure is from PDB ID: 5CCG. See **Supplementary Methods** for details.
[c] The Pearson and Spearman correlations between the smFRET-measured distances and the Cα-distances from the reference structure are 0.91 and 0.70, respectively.
[d] The first ranked pose predicted by DRSIP. See **Figure 5A** and **Table 1**.

**Table S3. MscL's smFRET data.**

| Residue Pair (*E. coli*)[a] | Residue Pair (*M. tuberculosis*)[a] | smFRET Dist. (nm)[b,d] | Ref Structure Dist. (nm)[c,d] | DRSIP Pred. Pose Dist. (nm)[e] |
|---|---|---|---|---|
| 25-25 | 23-23 | 4.6 | 1.2 | 1.2 |
| 27-27 | 25-25 | 4.5 | 0.8 | 0.7 |
| 42-42 | 40-40 | 5.0 | 2.0 | 1.9 |
| 75-75 | 69-69 | 5.2 | 2.4 | 2.3 |
| 80-80 | 74-74 | 4.4 | 2.1 | 1.9 |
| 82-82 | 76-76 | 5.0 | 2.1 | 2.0 |

[a] Residue pair are those between two neighboring monomers in the complex.
[b] smFRET-measured distances for the closed state of MscL computed from the experimental $R_0$'s and the FRET efficiencies (Wang *et al.*, 2014) by: $R_0 \sqrt[6]{\frac{1}{E} - 1}$.
[c] The reference structure is from PDB ID: 2OAR. Distances are between Cα atoms of the respective residue pairs.
[d] The Pearson and Spearman correlations between the smFRET-measured distances and the Cα-distances from the reference structure are 0.61 and 0.66, respectively.
[e] The second ranked pose predicted by DRSIP. See **Figure 5B and 5C** and **Table 1**.

#### Table S4. *Ct*H⁺-PPase and *Vr*H⁺-PPase smFRET data.

| **Residue Pair**[a] | | **smFRET Dist. (nm)**[b] | **Ref Structure Dist. (nm)**[c,d] | | **Docking Pose Dist (nm)**[e] | |
|---|---|---|---|---|---|---|
| ***Ct*H$^+$-PPase** | ***Vr*H$^+$-PPase** | | ***Ct*H$^+$-PPase** | ***Vr*H$^+$-PPase** | ***Ct*H$^+$-PPase** | ***Vr*H$^+$-PPase** |
| 170-170 | 243-243 | 4.23 | 3.71 | 3.72 | 3.92 | 3.59 |
| 219-219 | 292-292 | 5.19 | 5.13 | 5.14 | 5.17 | 5.08 |
| 411-411 | 499-499 | 4.24 | 5.49 | 5.50 | 5.45 | 5.51 |
| 452-452 | 540-540 | 4.20 | 4.02 | 4.03 | 3.95 | 4.02 |
| 642-642 | 736-736 | 4.22 | 2.89 | 2.90 | 3.00 | 2.83 |

[a] Residue pair are residues between 2 neighboring monomers in the complex. smFRET distances for *Vr*H$^+$-PPase are mapped from *Ct*H$^+$-PPase based on the pairwise sequence alignment (**Supplementary Fig. S3**) generated by SwissModel.
[b] smFRET-measured distances are from (Huang *et al.*, 2013).
[c] The reference structure for *Ct*H$^+$-PPase is the homology modelled structure (template structure: PDB ID: 4A01) and *Vr*H$^+$-PPase is PDB ID: 4A01. See **Supporting Methods** for details. Distances are between Cα atoms of the respective residue pairs.
[d] The Pearson and Spearman correlations between the smFRET-measured distances and the Cα-distances from the reference structure of *Ct*H$^+$-PPase and *Vr*H$^+$-PPase are 0.48 and 0.60, respectively.
[e] The first ranked pose for both systems predicted by DRSIP. See **Figure 5D and 5E** and **Table 1**.

**Table S5. Data of the three non-$C_n$ Symmetric HoTPs.**

| PDB ID | Num. of Chains | Chain 1 | Chain 2 | Dev. Ideal Rot Angle (°) | Shift Along Axis of Symm. (Å) | $C_n$-Symm. RMSD (Å) | Conform. RMSD (Å)[a] |
|---|---|---|---|---|---|---|---|
| **2GIF** | **3** | **A** | **C** | 6.18 | 0.34 | 3.38 | 1.47 |
| | | **B** | **A** | 2.46 | 2.04 | 2.48 | 1.09 |
| | | **C** | **B** | 3.65 | 2.49 | 3.26 | 1.72 |
| **2QTS** | **3** | **A** | **C** | 6.67 | 4.41 | 4.56 | 2.18 |
| | | **B** | **A** | 17.96 | 2.60 | 4.65 | 2.74 |
| | | **C** | **B** | 10.46 | 6.77 | 7.05 | 2.90 |
| **5HK1** | **3** | **A** | **B** | 1.30 | 21.13 | 21.15 | 1.76 |
| | | **B** | **C** | 41.59 | 24.45 | 33.44 | 1.48 |
| | | **C** | **A** | 35.85 | 10.36 | 31.17 | 0.91 |

[a] Conformational RMSD between the two chains.

#### Table S6. CAPRI criteria for ranking docking poses.

| Rank | % Nat | | iRMSD (Å) | | lRMSD (Å) | | Order of Symmetry (for HoTPs only) |
|---|---|---|---|---|---|---|---|
| High | ≥50 | and | (≤1.0 | or | ≤1.0) | and | (Identical to ref.) |
| Medium | ≥30 | and | (≤2.0 | or | ≤5.0) | and | (Identical to ref.) |
| Acceptable | ≥10 | and | (≤4.0 | or | ≤10.0) | and | (Identical to ref.) |
| Incorrect | <10 | or | (>4.0 | and | >10.0) | or | (Different from ref.) |

Reproduced with modifications from Méndez et al. (Méndez et al., 2003). The original CAPRI criteria do not consider the order of symmetry for HoTPs when evaluating the ranking. For evaluating HoTPs, the order of symmetry criterion is used, where the docking pose must have the same order of symmetry as the reference structure in addition to the CAPRI criteria. To obtain the rank of a docking pose, identify the highest rank such that the criteria is fulfilled by the pose. Take for example, a $C_3$ pose with % Nat: 30%, iRMSD: 3.0Å, lRMSD: 4.0Å and the reference structure is $C_3$, fulfills the criteria for the Acceptable and Medium ranks. The pose is assigned the Medium rank because it is the highest of the two ranks.

#### Table S7. Tilt Angle Criterion Reproduces the Size Distribution of $C_n$ Symmetric Poses in the HoTP Dataset (X-ray Structures).

| System | Type | Num. -mers | Chain Len. | $C_n$ Symm Criterion (Count & %)[a] | | | | | | $C_n$ Symm & Tilt Angle Criterion (Count & %)[b] | | | | | |
|---|---|---|---|---|---|---|---|---|---|---|---|---|---|---|---|
| | | | | $C_2$ | $C_3$ | $C_4$ | $C_5$ | $>C_5$ | Total | $C_2$ | $C_3$ | $C_4$ | $C_5$ | $>C_5$ | Total |
| **Pentameric Capsid Protein (4dmi)** | Soluble | 5 | 176 | 1507 | 104 | 160 | 145 | 203 | 2119 | | | | | | |
| **% Total** | | | | 71.1 | 4.9 | 7.6 | 6.8 | 9.6 | 100.0 | | | | | | |
| **MscL (2oar)** | Membrane | 5 | 174 | 1980 | 162 | 177 | 128 | 175 | 2622 | 298 | 25 | 56 | 39 | 50 | 468 |
| **% Total** | | | | 75.5 | 6.2 | 6.8 | 4.9 | 6.7 | 100.0 | 63.7 | 5.3 | 12.0 | 8.3 | 10.7 | 100.0 |
| **GH3 Beta-Glucosidase (5xxl)** | Soluble | 2 | 760 | 663 | 63 | 79 | 72 | 101 | 978 | | | | | | |
| **% Total** | | | | 67.8 | 6.4 | 8.1 | 7.4 | 10.3 | 100.0 | | | | | | |
| ***Vr*-PPase (4a01)** | Membrane | 2 | 766 | 826 | 95 | 40 | 45 | 61 | 1067 | 196 | 81 | 36 | 44 | 51 | 408 |
| **% Total** | | | | 77.4 | 8.9 | 3.7 | 4.2 | 5.7 | 100.0 | 48.0 | 19.9 | 8.8 | 10.8 | 12.5 | 100.0 |
| ***Ct*-PPase (SwissModel)** | Membrane | 2 | 673 | 723 | 134 | 53 | 47 | 98 | 1055 | 158 | 59 | 23 | 35 | 47 | 322 |
| **% Total** | | | | 68.5 | 12.7 | 5.0 | 4.5 | 9.3 | 100.0 | 49.1 | 18.3 | 7.1 | 10.9 | 14.6 | 100.0 |
| | | | | **$C_n$ Symm. HoTP Poses (Count & %)** | | | | | | | | | | | |
| **HoTP Dataset (X-ray Structures)** | Membrane | - | - | 47 | 26 | 25 | 8 | 9 | 115 | | | | | | |
| **% Total** | | | | 40.9 | 22.6 | 21.7 | 7.0 | 7.8 | 100.0 | | | | | | |

ZDOCK was used to dock and generate 54,000 docking poses for each system.

[a] The 54,000 poses were filtered with the $C_n$ symmetry criterion ($C_n$ RMSD > 2Å). The columns show the number of remaining poses and the percentage of the total for dimer, trimer, tetramer, pentamer and >pentamer.

[b] The 54,000 poses were filtered with the $C_n$ symmetry and tilt angle criteria (tilt angle > 35°).

### Supplementary Methods

#### Cyclic Symmetry-Imposed Packing (SIP)

##### *$C_n$ Symmetry RMSD Criterion*

The $C_n$ symmetry RMSD measures the deviation of the current pose from the ideal $C_n$ symmetric pose. This is used as part of the SIP filter to remove non-$C_n$ symmetric poses which are defined as poses with >2Å $C_n$ symmetry RMSD.

The $C_n$ symmetry RMSD is defined as

$$RMSD_{C_n} = \sqrt{\frac{\sum_{m=1}^{M} E_{1,m}^2 + E_{2,m}^2 + E_{3,m}^2}{M}} \tag{5}$$

where $E_{i,j}$ is the $i$'th row and $j$'th column element of the error matrix $\mathbf{E}_{3xM} = \mathbf{A'^*} - \mathbf{A'}$. **A'** and **A'$^*$** are coordinate matrices from the docking pose and ideal $C_n$ symmetric pose containing $M$ atoms.

There are two monomers in each pose where A is the first monomer and A' is the other. If there are no conformational differences between A and A', then A is used to build A'$^*$. Otherwise, A' is first superimposed onto A and then used to build A'$^*$ to remove the contribution of conformational differences from $C_n$ symmetry RMSD.

##### *Tilt Angle Criterion*

The membrane normal is assumed to be the axis of symmetry, $\hat{\mathbf{r}}$.

Each monomer is modeled as a cylinder spanning the membrane in the direction of the cylindrical axis ($\mathbf{c}_{axis}$) which is parallel to height of the cylinder. $\mathbf{c}_{axis}$ is the sum of the longitudinal axes (unit vectors) of the transmembrane helices (Cα atoms only) in the monomer. Principal component analysis (PCA) (Yang *et al.*, 2009; Chandrasekaran *et al.*, 2016) is used to identify the longitudinal axis (eigenvector) having the largest variance (the corresponding eigenvalue) of the atom's positions with respect to the COM of the helix.

The tilt angle of a monomer, $\theta_t$, is defined as the acute angle between the rotation axis $\hat{\mathbf{r}}$ and $\mathbf{c}_{axis}$. This criterion checks if one of the monomers in the docking pose has $\theta_t \leq 35°$.

The residues that form the transmembrane helices are assigned using the method proposed by Lomize et al. (Lomize *et al.*, 2011).

#### Distance-Restraints (DR) Filter

Pearson and Spearman correlation coefficients are used to evaluate the goodness of fit between the smFRET data and the corresponding distances in docking poses. Docking poses with ≤0.3 correlation as measured by Pearson or Spearman correlation are filtered out.

#### Shift Along Axis of Symmetry

Shift along axis of symmetry is defined as $\left|\mathbf{d}_{diff}^T \hat{\mathbf{r}}\right|$ (from **Equation (4)**), which is the absolute value of the projection of the difference vector ($\mathbf{d}_{diff}$) onto the axis of symmetry ($\hat{\mathbf{r}}$) where $\hat{\mathbf{r}}$ is a unit vector.

#### Deviation from Ideal Rotation Angle

The deviation from ideal rotation angle is defined as $|\theta_n - \theta_r|$ where $\theta_r$ is the angle of rotation obtained from the $C_n$ symmetry rotation matrix (**R** in **Equation (1)**) while $\theta_n = \text{sign}(\theta_r)\frac{2\pi}{n}$ is the ideal rotation angle with $n = |2\pi/\theta_r|$ rounded to the nearest integer.

#### Clustering of Poses

The remaining docking poses after applying the filters were clustered using agglomerative clustering with the average linkage criteria (Manning *et al.*, 2008). In agglomerative clustering, a pair of clusters with the shortest distance are combined into a new cluster and the distance between this new cluster with the other clusters are recomputed. This is done iteratively until the shortest distance is more than the cutoff distance. While, the average linkage criteria define the distance between any two clusters as the average distance between all pairs of members, one from each cluster.

RMSD is used to quantify the difference/distance between all pairs of poses. The cutoff is 12 Å and clusters with fewer than three poses are removed considering the disfavored entropy.

To calculate the RMSD, for each pair of poses (each pose contains a dimer), the two dimers are first superimposed at one monomer taken from each dimer and then the RMSD between the remaining monomers are computed. There are four possible combinations of monomers that can be chosen for the superimposition and the one with the lowest RMSD is chosen to represent the distance between the two poses.

##### *Selecting the Representative for Each Cluster and Ranking of the Clusters*

The clusters for both HoTPs and soluble proteins are rank-ordered based on the size of the clusters so that the first cluster has the most members. The representative pose of each cluster is then chosen by the following rules for HoTPs and soluble proteins.

Within each cluster of HoTPs, the consensus order of symmetry is determined by a simple majority. Among the members of the cluster who has the consensus order of symmetry, the pose that has the lowest $C_n$ Symmetry RMSD is chosen as the representative of the cluster.

The representative of a cluster for soluble proteins is chosen by selecting the member in the cluster with the highest distance restraints' Spearman correlation. If there are multiple members with the same highest Spearman correlation, then among the members, the one with the highest distance restraints' Pearson correlation is selected as the representative.

#### Data Sets

##### *Curation of Non-Redundant X-ray Solved α-helical HoTP Data Set*

The PDB IDs of α-helical transmembrane proteins are from the mpstruct database (http://blanco.biomol.uci.edu/mpstruc/). The PDB IDs were filtered using the advanced filter of RCSB PDB (Rose *et al.*, 2017) with the following criteria: X-ray structures have ≤4Å resolution, biological assembly is a homo-oligomer, transmembrane protein is α-helical and <30% sequence identity between any pair of sequences, returning 152 structures.

Among the 152 structures, 129 were manually verified to be transmembrane proteins and homo-oligomers. Extra-membrane domains >30 residues were removed from the structures. The domains were determined based on the annotations of MemProtMD (Stansfeld *et al.*, 2015) and OPM (Lomize *et al.*, 2011).

#### *Dihedral Symmetric Structures in HoTP Data Set*

Dihedral symmetric HoTP structures ($D_n$) contain two complexes and two orthogonal axes of symmetries, one that is the membrane normal and other that is parallel to the membrane bilayer. Each $D_n$ structure contains two identical $C_n$ symmetric complexes whose axes of symmetry is parallel to the membrane normal. The two complexes are related by a $C_2$ symmetry about the axis parallel to the membrane normal (Ghelis and Yon, 1982).

Two dihedral symmetric structures are included in the 118 parallel HoTPs by taking one of the two complexes.

#### *Experimental Data Used for Validating smFIRST*

The Syt1-SNARE smFRET data are from Choi et al (Choi *et al.*, 2010). Only smFRET pairs that have a single Gaussian peak and the peak is not too broad such that the difference between the maximum and minimum FRET efficiency that the peak covers is ≤50%, are included (**Supplementary Fig. S1**, **Table S1 and S2**). The reference structure for the Syt1-SNARE complex is PDB ID: 5CCG's complex 1 - interface 1 (Zhou *et al.*, 2015). The Syt1's GLU140 which is one of the smFRET pairs is missing. Therefore, the structure of Syt1 was taken from PDB ID: 2R83 (Huang *et al.*, 2013) but the relative orientation of the two domains (C2A and C2B) in Syt1 are different from the reference structure. Each domain was individually superimposed onto the corresponding domain in the reference structure. The RMSD difference between the Syt1 in the reference structure with the Syt1 created from superimposing the two domains is 1.21 Å.

For MscL, the smFRET data (**Supplementary Table S3**) are for the closed state *E. coli* MscL (Wang *et al.*, 2014). The smFRET distances are calculated from the smFRET efficiencies and $R_0$ provided by the authors. The reference structure of MscL (PDB ID: 2OAR) is from *M. tuberculosis* which is different from the smFRET data therefore multiple sequence alignment (**Supplementary Fig. S2**) between the MscL sequences from *M. tuberculosis*, *E. coli*, *M. leprae*, *B. subtilis* and *H. influenzae* were performed with MUSCLE (Edgar, 2004; McWilliam *et al.*, 2013). The smFRET residue pairs from the *E. coli* MscL were then mapped to those in *M. tuberculosis* MscL according to the alignment (**Supplementary Fig. S2** and **Supplementary Table S3**). Chains A and B from the reference structure are used to evaluate the quality of the docking poses. Chain A was replicated and the two replicas were docked using ZDOCK.

The structure of the *Ct*$H^+$-PPase system was predicted by homology modeling using SwissModel (Biasini et al., 2014) with the template structure (PDB ID: 4A01) obtained from the *Vr*$H^+$-PPase system (Sequence identity: 45.1%, **Supplementary Fig. S3** and **Table S4**). The predicted structure is $C_2$ symmetric and used as the reference structure. Chain A was replicated and used to perform docking. While, the smFRET data for *Ct*$H^+$-PPase bound to $K^+$ and Mg-IDP is from Huang et al. (Huang *et al.*, 2013).

As for *Vr*$H^+$-PPase, the *Ct*$H^+$-PPase smFRET data are mapped to the corresponding residue pairs in *Vr*$H^+$-PPase based on the pairwise sequence alignment (**Supplementary Fig. S4** and **Table S4**) generated by SwissModel. The reference structure for *Vr*$H^+$-PPase is PDB ID: 4A01.

#### Quality of Docking Poses: CAPRI Criteria

Critical Assessment of PRediction of Interactions (CAPRI) is a community project that performs blind experiments to evaluate the performance of current methods for predicting protein-protein interactions.

The CAPRI criteria are used to evaluate the quality of the docking poses with respect to the native pose from X-ray structures. The CAPRI criteria (see **Supplementary Table S6**) uses percent native contacts (%Nat), interface RMSD (iRMSD) and ligand RMSD (lRMSD) to classify the poses into one of the following ranks: Incorrect, Acceptable, Medium or High (Méndez *et al.*, 2003). Native poses are defined as a pose that is not "Incorrect".

Native contacts are residues from the two monomers in the native pose whose heavy atoms are within 5Å of one another. The %Nat is the percent of contacts in the native pose that exists in the docking pose. Since the monomers are identical and there are two binding interfaces on each monomer in $C_n$ symmetric poses, there are two ways to compute %Nat and the highest is used. For the first way to compute %Nat, all the residues involved in the native contacts on the first monomer of the native pose are taken from the static pose while the residues from the second monomer are taken from the mobile pose for computing the contacts of the docking pose. The other way to compute %Nat, the residues taken from the two monomers in the docking pose are swapped. The order of the static and mobile monomers which results in the higher %Nat is used for computing iRMSD.

iRMSD is the RMSD of the backbone atoms computed between the interface residues from the docking pose and the reference structure. The interface residues are defined as those with heavy atoms within 10Å between the two monomers in the reference structure. Before calculating the iRMSD, the docking pose and native structure are superimposed at the binding interface.

The ligand RMSD (lRMSD) takes one of the two identical monomers in a pose as the "receptor" and is used to superimpose the docking and the native poses. After superimposition, the RMSD of backbone atoms between the "ligands" (monomers that were not used in the superimposition) are calculated. Understandably, there are four possible ligand-receptor combinations. The combination that gives the lowest lRMSD is chosen.

#### Selection of Soluble Proteins for Comparison to MscL and *Vr*$H^+$-PPase/*Ct*$H^+$-PPase

Two soluble proteins were chosen such that they are of similar sizes to their HoTP counterparts. This is achieved by filtering the RCSB PDB database such that the protein is soluble, $C_5$ ($C_2$) symmetric and whose chain length is within +/- 10 residues of MscL (*Vr*$H^+$-PPase), which is between 164 to 184 (756 to 776) residues long. From the filtered results, the Pentameric Capsid

Protein (PDB ID: 4dmi) and GH3 Beta-Glucosidase (PDB ID: 5xxl) were chosen for comparison to MscL and *Vr*$H^+$-PPase/*Ct*$H^+$-PPase, respectively. See **Table S7**.

### Supplementary Results

#### Anti-Parallel HoTPs

Among the 129 HoTPs in the dataset, 11 of them are anti-parallel. Out of the 11 structures, 8 of the anti-parallel HoTPs have not been verified through other experiments or not implicated in the proper functioning of these proteins. It is possible that these anti-parallel HoTPs are due to crystal contacts rather than real interactions under physiological conditions. However, three proteins are identified to be *de facto* anti-parallel.

The first is the dimeric multidrug transporter from *Escherichia coli*, EmrE (PDB ID: 3B5D) which has X-ray, cryo-EM and smFRET experimental evidence supporting the anti-parallel orientation (Du et al., 2015).

While the dimeric de novo designed $Zn^{2+}$ transporter (PDB ID: 4P6J) has experimental data from NMR validating the anti-parallel orientation observed in the X-ray structure (Joh et al., 2014).

Finally, the anti-microbial peptide channel dermcidin (PDB ID: 2YMK) which is a $D_3$ hexamer, where each monomer in the complex spans the same membrane bilayer, is required to form a channel (Song et al., 2013).

#### The Three Non-$C_n$ Symmetric Structures

The three non-$C_n$ symmetric HoTPs (**Supplementary Table S5**) among the 118 parallel HoTPs that have $C_n$ symmetry RMSDs >2Å (cutoff) are the human $\sigma_1$ receptor protein (PDB ID: 5HK1), acid-sensing ion channel protein (PDB ID: 2QTS) and major drug-resistance pump AcrB (PDB ID: 2GIF).

The first protein, the human $\sigma_1$ receptor (PDB ID: 5HK1, **Supplementary Fig. S4A**) is not a proper HoTP because its transmembrane region acts as a membrane anchor but the transmembrane regions do not interact with one another in the membrane (Schmidt *et al.*, 2016).

While the acid-sensing ion channel protein (PDB ID: 2QTS, **Supplementary Fig. S4B**) is expressed without its native N- and C- terminal residues and is functionally inactive (Jasti *et al.*, 2007). It is likely that the truncated termini disrupted the structure and function of the protein.

Finally, the trimeric protein major drug-resistance pump AcrB (PDB ID: 2GIF, **Supplementary Fig. S4C**) where each monomer is suggested by Seeger to represent the "consecutive states of a transport cycle" and that the protonation state of titratable groups in each monomer are different (Seeger *et al.*, 2006). The different protonation states of each monomer probably effects the way each monomer interacts with its neighboring monomers thus disrupting the $C_n$ symmetry of the complex.
